# IL-2 secreting T helper cells promote EF B cell maturation via intrinsic regulation of B cell mTOR/AKT/Blimp-1 axis

**DOI:** 10.1101/2022.08.19.504576

**Authors:** Caterina E. Faliti, Maria Mesina, Jinyong Choi, Simon Bélanger, William R. Schief, Shane Crotty

## Abstract

**SUMMARY:** B cells are fundamental players in the secretion of antibodies and the establishment of long-term memory-based immunity. Integration of signals from TLRs, BCR stimulation, and T helper cell-derived cytokines can all dictate B cell differentiation and their metabolic state. However, while important components of this interaction have been described, the precise signaling networks and mechanisms regulating B cell fate are not fully understood. Here, we elucidated the role of interleukin-2 (IL-2) in determining early B cell fate decisions and inducing plasma cell reprogramming. Using both *in vitro* culture systems and *in vivo* models of immunization, alongside CRISPR-based genome editing of antigen-specific T and B cells, we identify a role for T helper-secreted IL-2 in inducing high expression of Irf4 and Blimp-1 in activated cognate B cells, enhancing plasma cell differentiation. Induction of this cascade promotes their differentiation and drives metabolic reprogramming through the regulation of mTOR/AKT/Blimp-1 axis.

## INTRODUCTION

Plasma cells are terminally differentiated B cells that can secrete large amounts of protective antibodies during infections or vaccination 8/19/22 11:12:00 AM. Antibodies are the first line of defense against pathogens beginning in the acute phase of infection and persisting beyond recovery through homeostatic long-term memory formation and the generation of long-lived plasma cells (LLPC). Vaccine-induced immunity and protection against pathogens ultimately rely on the induction of antigen-specific long-lived antibody-secreting cells capable of clearing the pathogen *in vivo* (Slifka et al.,1998). Despite their importance in pathogen control and vaccination, the factors that influence B cell fate determination and plasma cell formation have yet to be fully understood.

Upon naïve B cells activation, canonical responses are characterized by early induction of extrafollicular (EF) B cells populations of low-mutated plasmablasts (EF B_PB_) and early minted plasma cells (EF B_PC_) followed by sequential rounds of divisions and maturation of selected B cells inside the germinal centers (GCs) (Elsner & Shlomchik, 2020). GCs are microanatomical structures normally present in secondary lymphoid organs where T cell interactions and repeated B cell receptor (BCR) testing drive affinity-matured class-switched GC B cells (B_GC_). Early in this process, activated antigen-specific B cells interact with cognate T cells at the T-B border, where they compete for T cell help. Based on their frequency, affinity, or relative access to the antigen (Woodruff et al., 2018), B cells can proceed to enter a GC, where they can undergo several rounds of proliferation, class-switch recombination (CSR) and somatic hypermutation (SHM) (Eisen & Siskind, 1964; Victora et al., 2012). Later GCs outputs are high-affinity memory B cells and bone marrow resident LLPC.

The dominance of EF versus GC-mediated processes varies according to the system investigated, with non-canonical responses largely observed in the context of infections (Di Niro et al., 2015) or autoimmunity (Jenks et al., 2018), where activated naïve B cells serve as a continuous supply for the generation of B_PB/PC_ at the EF sites. EF sites are in fact critical in certain aspects of humoral immunity, where both T-independent and T-dependent mechanisms can instruct blasting B cells to differentiate into antibody-secreting cells (ASC) capable of undergoing CSR and persist for a longer period (Jenks et al., 2019; Roco et al., 2019).

Sequential integration of three signals contributes to the differentiation of naïve B cells: BCR/antigen cross-linking, toll-like receptors (TLRs) activation on their surface, and cognate interaction with T helper cells (Akkaya et al., 2018; Ruprecht & Lanzavecchia, 2006). Upon antigenic stimulation, B cells rapidly increase their metabolic activity and prepare to receive signals from TLR ligation and cognate T cells interaction. Activated B cells express a variety of TLRs which can participate in the shaping of B cells responses and their differentiation (Hua & Hou, 2013). TLR4, activated by LPS, is almost uniquely expressed on mouse B cells (Buchta & Bishop, 2014), while TLR9, activated by CpG, is widely expressed on both mouse and human B cells. Therefore, TLR9 represents a valuable target to study for understanding B cell biology. CpG used to activate TLR9 in the form of an adjuvant has had relevant outcomes both *in vitro* and *in vivo* (Akkaya, Akkaya, Miozzo, et al., 2017; Akkaya, Akkaya, Sheehan, et al., 2017; Coffman et al., 2010; Sanjuan Nandin et al., 2017).

T cell help is provided in a dynamic, multifactorial process in which activated T helper cells receive sustained stimulation from and can provide costimulatory and survival signals, as well as secreted cytokines to B cells ultimately favoring their differentiation (Crotty, 2015). Classical T help provided to B cells consists of IL-21, IL-4, CD40L, and CXCL13. Additionally, IL-2 and TNF cytokines have also been implicated in B cell maturation processes (Crotty, 2015, 2019).

IL-2 is a pleiotropic cytokine discovered in the early 1980s (Robb & Smith, 1981) able to bind with high affinity to the three chains of the IL-2 receptor, termed alpha (α, CD25), beta (β, CD122), and gamma (γ, CD132) (Smith, 2006). IL-2 biology on T cells has been extensively investigated, with fundamental understanding achieved for how IL-2 mediates T cell survival, proliferation, cell fate differentiation, and metabolism. In T cells, IL-2 activity has been linked to cell fate regulation of Th1/Tfh (DiToro et al., 2018) and Blimp-1-dependent mTOR activation (Ray et al., 2015). Blimp-1, a master gene regulator involved in B_PC_ differentiation and LLPC maintenance (Shapiro-Shelef et al., 2005), has also been implicated in fueling energetic requirements during metabolic changes and restructuring of B cells that allow for the secretion of high amounts of glycosylated antibodies (Boothby & Rickert, 2017).

Here, we investigated the precise role of IL-2 cytokine secreted by T helper cells during their cognate interactions with activated B cells. Adopting a mouse model of immunization with HIV-related immunogens, we explored this relationship using CRISPR-based genome editing of antigen-specific T and B lymphocytes, efficiently generating knockout cells insensitive to IL-2 signaling and viable for further *in vitro* and *in vivo* investigations. Using this approach, we identified the participation of IL-2 signaling early in B cell activation guiding both metabolic restructuring and plasma cell differentiation. Mechanistically, we observed that IL-2 acts intrinsically on B cells to promote the expression of Irf4 and Blimp-1 in a BACH2-independent manner. The induced expression of Blimp-1, specifically, is key to promoting the mTOR-linked signaling required for B_PC_ terminal differentiation.

## RESULTS

### IL-2 increases EF B_PC_ differentiation and antibody secretion during i*n vivo* immunization

Cytokines are potent mediators of immune cell activation and effector functions. They have pleiotropic effects and can participate in several aspects of immune responses in both health and disease conditions (Dinarello, 2007). While the functions of key T helper secreted cytokines, such as IL-21 and IL-4 have been well characterized (Bélanger & Crotty, 2016), the precise role for IL-2 in mediating T-B cell interactions remains unclear. The wide secretion of IL-2 upon T cell engagement with the antigen and the critical role of activated T helper cells in governing B cell differentiation warrants deep investigation of the role of IL-2 in shaping adaptive immunity and supporting immune-modulation.

To elucidate the role of IL-2 in tuning adaptive immune responses upon *in vivo* immunization, an antigen-specific model designed to track cognate T cells and B cells following vaccination was developed. Simultaneous adoptive transfer of transgenic VRC01^gHL^ uGFP B cells and Smarta CD4 T cells into congenic C57BL/6 recipient mice followed by immunization with an alum-adsorbed eOD-GT5_gp61_60-mer immunogen allowed antigen-specific tracking throughout development of humoral immunity. As previously described (Kato et al., 2020), the incorporation of the “gp61-80” LCMV Smarta peptide “GLKGPDIYKGVYQFKSVEFD” into the design of the eOD-GT5 60-mer immunogen links the Smarta cells and VRC01^gHL^ B cell responses *in vivo*.

Previous studies have identified the differentiation of antigen reactive VRC01^gHL^ B cells along with the EF and GC trajectories as influenced by several factors including precursor frequency (Abbott et al., 2018), affinity and avidity for the immunogen (Kato et al., 2020), and the cognate T cell help received (Lee et al., 2021). Since the initial input of transferred precursor naïve VRC01^gHL^ B cell could influence their fitness and differentiation *in vivo* (Abbott et al., 2018), we tested both low (1 in 10^6^) and high (1 in 10^4^) numbers of precursor cells transferred into recipient mice.

Immunized cohorts received a daily intravenous injection of recombinant IL-2 (rIL-2) starting three days post-immunization, thus allowing initial pre-Tfh cell differentiation to develop mostly unperturbed (Y. S. Choi et al., 2011) (Figure 1A). One-week post-immunization, spleens of rIL-2 treated mice showed increased splenocyte counts (Figure S1A), with increasing counts of both T and B lymphocytes (Figure S1B and S1E). Among CD4 T cells, the frequency of Tfh cells was unaltered, although the frequency of Smarta Tfh cells was reduced – likely due to sustained Blimp-1 expression induced in these cells by IL-2 (Figure S1C). At this time of analysis, conversely to influenza infection models (Botta et al., 2017) we did not observe a modulation of Foxp3-expressing T cells upon IL-2 injections (Figure S1D). Absolute B cell counts were also increased upon IL-2 treatment (Figure S1E). Assessment of B cell fate decisions revealed that the IL-2 treated group displayed significant expansions of EF B_PB/PC_ over B_GC_ when comparing frequencies for both endogenous (Figure 1B-1D) and VRC01^gHL^ B cells (Figure 1E-1H), while EF B_PB/PC_, B_GC_, and VRC01^gHL^ B_GC_ cell absolute numbers were all increased (Figure S1F-1G).

**Figure 1.**
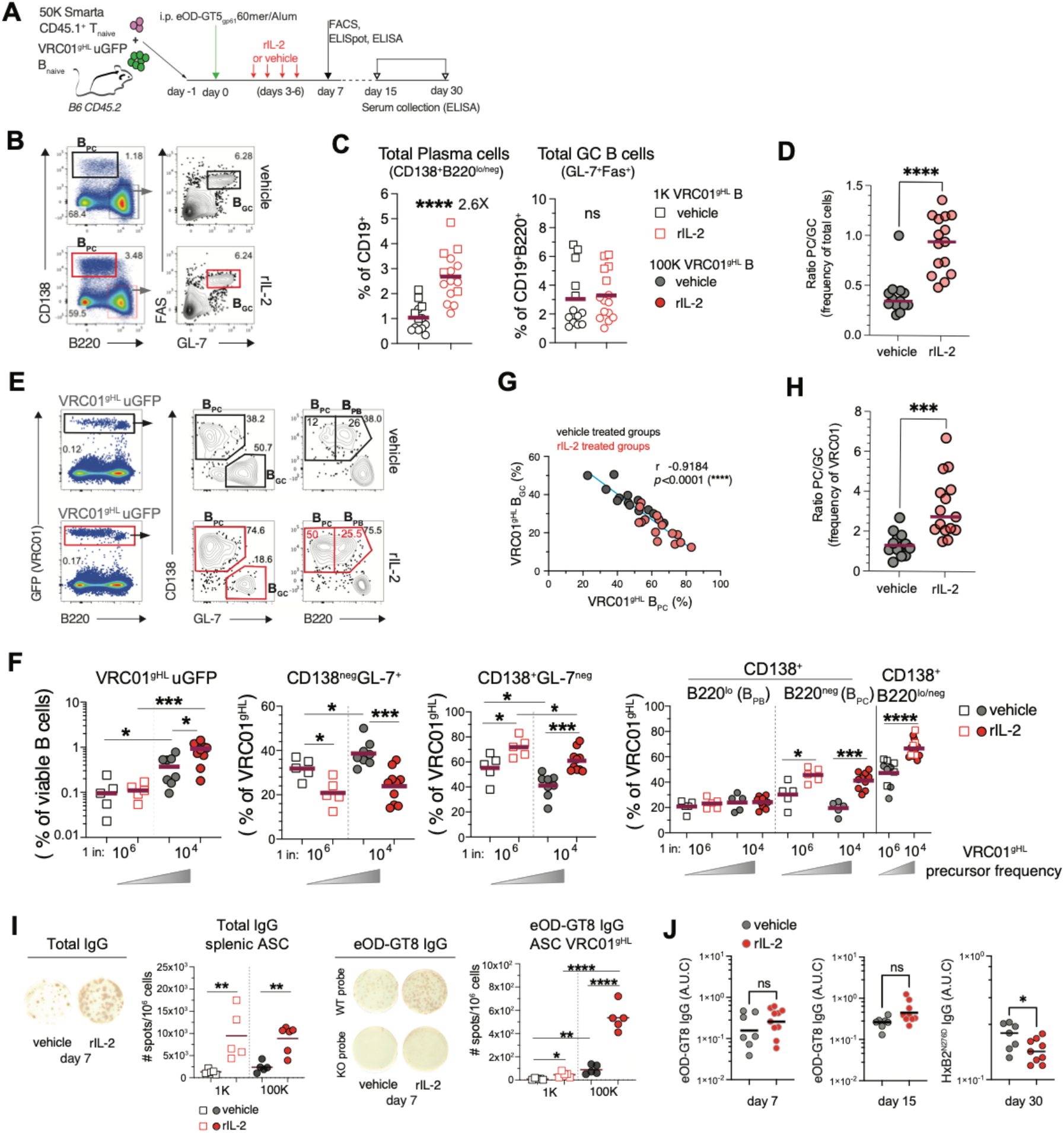
In vivo IL-2 administration enhances plasma cells differentiation and antibody secretion upon eOD-GT5 immunization. (A)Schematic representation for adoptive cell transfer of Smarta CD45.1+ T cells (number of cells transferred: 50K) and VRC01^gHL^ homo uGFP B naïve cells (number of cells transferred: 1K (squares in graphs) or 100K (circles in graphs)) into B6 CD45.2^+^ recipient mice, a day prior intra-peritoneally (i.p.) immunization with 20 µg eODGT5_gp61_60mer precipitated in Alum. Mice were treated with r-IL2 (50K U) or vehicle (PBS) administered daily via retro-orbital intravenous (i.v.) injection in 50 µl, from day 3 to 6 post-immunization. Analysis on spleens performed as indicated, and serological assessment to quantify antigen-specific IgG. (B) Representative dot plots for the gating of total PC and GC B cells in spleens and (C) their frequencies assessed in the groups of vehicle controls and rIL-2 treated mice. (D) Ratio for PC over GC frequency of total B cells. (E) FACS plots showing the gating strategy to quantify the VRC01^gHL^ GFP^+^B220^+/-^ B cells gated on viable, CD3^−^CD19^+^ cells. VRC01^gHL^ homo uGFP cells were scored for their differentiation into GC (GL7^+^CD138^−^), PC (GL7^−^CD138^+^ or B220^−^CD138^+^) and PB (B220^+/lo^ CD138^+^). (F) Quantifications of transferred VRC01^gHL^ from different precursor frequencies (pf) (low pf=1K, 10^6^ and high pf=100K, 10^4^) are shown for each population. (G) Correlation between VRC01 GC and PC in the two cohorts of analyzed mice (H) Ratio for PC over GC frequency of VRC01^gHL^ B cells. (I) Enumeration of IgG secreting ASC from splenocytes isolated on day 7 and tested in ELISpot for the detection of total and antigen-specific eOD-GT8 IgG. (J) Serological evaluation of circulating antibodies and affinity maturation over time from sera of all immunized controls or rIL-2 treated mice tested in ELISA against eOD-GT8 and HxB2^N276D^ probes. Each dot represents an individual mouse. Lines in graphs represent the mean value. Data are combined from n=2 experiments for each group of mice transferred with low or high pf. P values were determined by two-tailed Student’s t test. Not significant, ns p > 0.05; *p < 0.05; **p < 0.01; ***p < 0.001; ****p < 0.0001.

Previous literature has reported increased CD25 expression on lymphocytes as a measurement of their responsiveness to IL-2. In our model, we detected the highest levels of CD25 on endogenous CD4 T cells and EF B_PB/PC_ (Figure S1H-S1J). The observed heightened commitment into EF B_PB/PC_ differentiation resulted in a functional enhanced ability to secrete total and antigen-specific IgG (Figure 1I).

Priming of the transferred VRC01-class precursor B cells with the eOD-GT8 60mer promotes their clonal expansion and selects for mutations that can confer cross-reactivity to more native-like gp120 constructs, such as the core-e-2CC HxB2 N276D (HxB2^N276D^) a conformationally stabilized core gp120 monomer with a near-native CD4bs from strain HxB2 (Jardine et al., 2015). Serological evaluation of the circulating IgG induced upon immunization that bind to the germline-targeting eOD-GT8 or the CD4bs-like HxB2^N276D^ probes can be informative of the maturation and mutational states of the *in vivo* activated VRC01^gHL^ B cells. In our model, while the accumulation of circulating eOD-GT8 specific IgG was similar between the two groups, the level of HxB2^N276D^-reactive IgG was reduced in the mice receiving IL-2 injections (Figure 1J and S1K) suggesting that rewiring early IL-2 signals during immunization may imprint differentiation choices on activated B cells preferentially towards EF B_PB/PC_.

### IL-2 secreting Smarta T helper cells control the magnitude of VRC01^gHL^ plasma cells generated upon *in vivo* immunization

While establishing the importance of IL-2 in B cell response development, the immunization model has two main limitations: 1) the inability to define the cellular source of IL-2 and 2) the inability to understand if B cells can be intrinsically regulated through IL-2 signaling – a mechanism that has been largely excluded or considered indirectly dependent on T cells regulation (Ballesteros-Tato et al., 2012); Willerford et al., 1995).

As both mouse and human CD4 T cells share the ability to secrete IL-2 in response to antigenic stimulation (Mosmann et al., 1986; Whyte et al., 2020) and are largely responsible for its secretion in lymphoid tissues such as mouse spleen or lymph nodes (Whyte et al., 2020), they seemed the most likely source of soluble IL-2. Nonetheless, a detailed quantification and a direct comparison of IL-2 producing T helper subsets among CD4 T cells upon antigen stimulation has lagged. Therefore, to interrogate the relevance of IL-2 secreting T helper cells in human biology, we quantified the frequency of IL-2 secreting antigen-specific human T helper memory subsets circulating in blood or resident in tissues (Figure S2A and S2B). Cells were either incubated with “mega pools” (MPs) of peptides specific for the CMV or PT human pathogens or polyclonally stimulated *in vitro*. Intracellular cytokine secretion (ICS) assay allowed us to identify cells secreting IL-2, IFNγ and TNFα cytokines and expressing CD40L (as a measure of help provided to CD40^+^ B cells). We detected antigen-driven IL-2 secretion in memory T cells in ~40-90% of the healthy donors’ sample tested. Those CD40L^+^IL-2^+^ T cells were equally distributed between the CXCR5^+^ and CXCR5^−^ compartments in a range varying from 0.01-0.1% of total CD4s T cell subsets for antigen-specific cells (Figure S3A-S3D) and ~2-60% of polyclonal stimulated cells (Figure S3E and S3F). Co-secretion of IFNγ and/or TNFα varied within each tested stimuli, with CD40L^+^IL-2^+^ CXCR5^−^ cells displaying a slightly higher enrichment of those cytokines compared to their CXCR5^+^ counterparts (Figure S3G-S3H).

Similarly, mouse studies from our group (J. Choi et al., 2020) and others (DiToro et al., 2018) have shown that CXCR5^+^ Tfh are capable of secreting IL-2 *in vivo* (Figure S4A), suggesting that CD40L^+^IL-2^+^ T_FH_ cells are capable of shaping antigen-specific B cells differentiation during antigen-mediated immune responses in both mice and humans.

To assess whether IL-2-secreting Smarta T helper cells were responsible for controlling VRC01^gHL^ B cell differentiation in our model, we selectively disrupted the *Il2* locus in T cells with a CRISPR/Cas9-mediated retroviral editing approach and generated IL-2 disrupted (KO) cells (gIl2 Smarta) (Figure S4B and S4C). This method allowed us to obtain viable cells with high levels of transduction and gene KO efficiency (Figure S4D and S4E), as previously described (J. Choi et al., 2020).

IL-2 KO cells, or CD8a-knockout controls, were transferred into B6 recipient mice four days prior to immunization. One day prior to immunization, VRC01^gHL^ uGFP B cells were also transferred (Figure 2A). After one week, analysis of *ex-vivo* isolated gIl2 Smarta cells in comparison to gCd8a Smarta showed a similar differentiation into Tfh cells (Figure 2B and 2C) as well as similar levels of activation and differentiation (Figure S4F). Il2 disruption was stable in *ex-vivo* restimulated gIl2 Smarta cells on day 7 post-immunization (Figure 2D and 2E). Akin to findings with human CD4 T helper cells, both CXCR5^−^ and CXCR5^+^ CD40L expressing gCd8a Smarta cells were able to secrete IL-2 when restimulated *in vitro* (Figure 2F and 2G).

**Figure 2.**
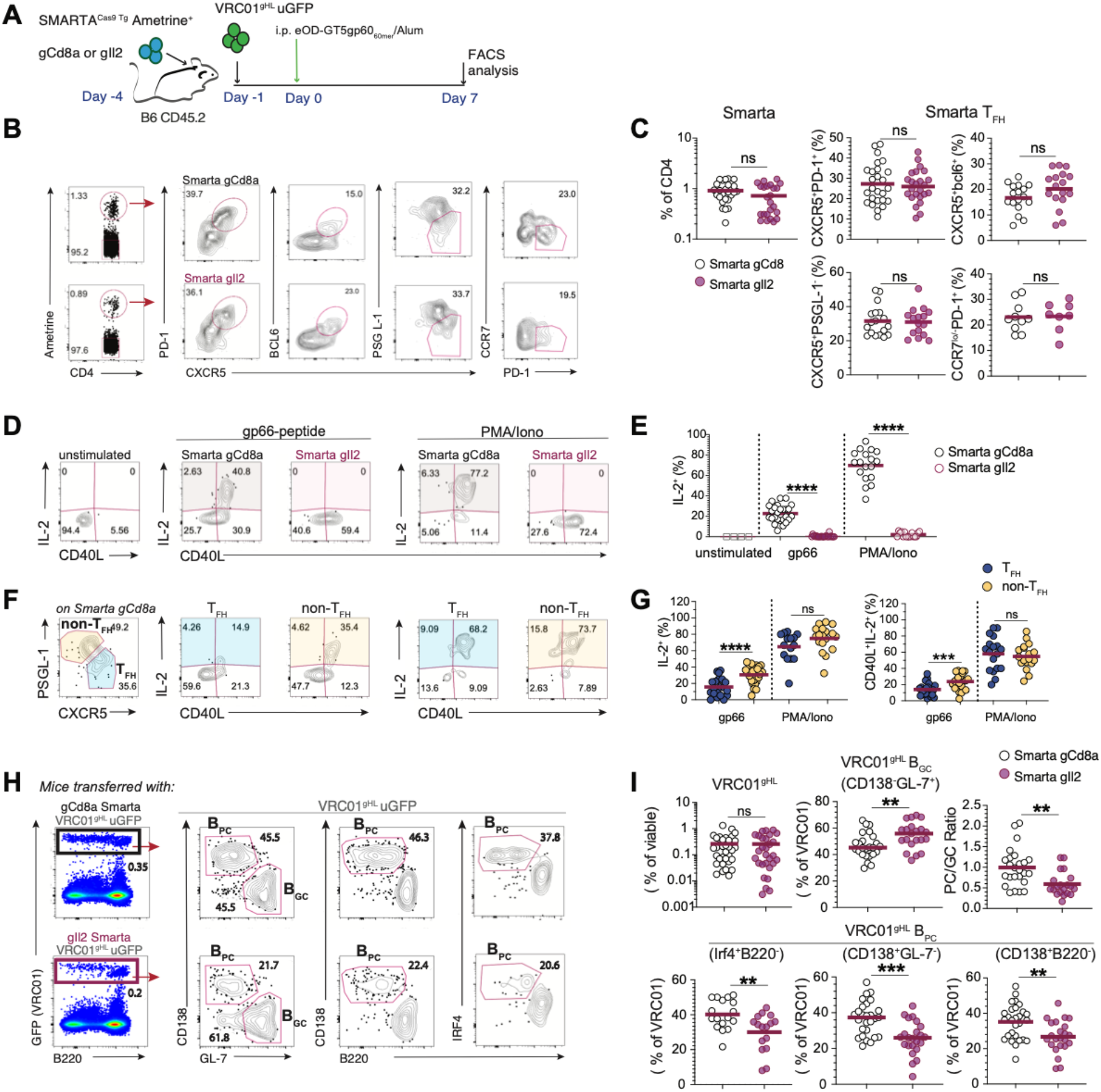
IL-2^+^ Smarta cells regulate VRC01^gHL^ cell GC or PC differentiation. (A)Schematic representation of adoptive cell transfer of CRISPR/Cas9 modified Smarta cells (gCd8 target cells as irrelevant guide control or gIl2 from IL-2 knock-out Smarta cells) and VRC01^gHL^ uGFP naïve B cells into CD45.2^+^ B6 recipient mice. Intra-peritoneal immunization was performed with administration of 20 µg of eODGT5^gp61^60mer immunogen precipitated in Alum. Spleens were analyzed after one week. (B) Representative dot plots of gRNA Smarta cells (CD4^+^Ametrine^+^) isolated from spleens on day 7 show their phenotype as T follicular helper cells based on the expression of bcl6, CXCR5, PD-1, PSGL-1, and CCR7. (C) Quantification of total Smarta and Smarta Tfh cells from mice recipient of gCD8 Smarta (controls, n=26) and gIl2 Smarta (KO, n=24). (D) ICS in vitro assay to quantify IL-2 secretion from Smarta cells. Splenocytes isolated on day 7 post-immunization were stimulated *in vitro* with either their cognate peptide gp-66 or PMA/ionomycin and analyzed at ~4 hours after incubation. FACS plots show IL-2 and the co-expression of CD40L on stimulated cells. (E) Quantification of IL-2 upon stimulation of gCd8 or gil2 Smarta. (F) FACS plots showing the gating strategy to identify gCd8 Smarta Tfh (CXCR5^+^PSGL-1^−^) and non-Tfh (CXCR5^−^PSGL-1^+^) subsets and expression of IL-2, CD40L. (G) Relative quantification of IL-2 secreting cells from each indicated population as in (F). Each dot represents an individual mouse. Data are combined from n=3 experiments. (H) FACS plots showing the quantification of VRC01^gHL^ cells (gated as viable, CD3^−^CD19^+^ GFP^+^B220^+/-^ B cells) and their differentiation into GC (GL7^+^CD138^−^) or PC (GL7^−^CD138^+^, B220^−^ CD138^+^ and B220^−^IRF-4^+^) scored on day 7. (I) Relative quantifications of each indicated population of VRC01^gHL^ B cells. Each dot represents an individual mouse. Lines in graphs represent the mean value. Data are combined from n= 3 or 4 experiments. P values were determined by a two-tailed Student’s t-test. Not significant, ns p > 0.05; *p < 0.05; **p < 0.01; ***p < 0.001; ****p < 0.0001.

To precisely interrogate the *in vivo* helper functions of IL-2 KO Smarta T cells, antigen-specific B cell differentiation was also analyzed one-week post-immunization. While the expansion of VRC01^gHL^ uGFP B cells was not different between the two groups of mice (Figure S4G), in the absence of IL-2 secreting Smarta cells their differentiation into B220^−^CD138^+^IRF4^+/hi^ B_PC_ was significantly reduced, in favor of an increased expansion of CD138^−^GL-7^+^ B_GC_ (Figure 2H and 2I).

Taken together, these findings demonstrate that IL-2 secreting Smarta T helper cells are required to effectively imprint on VRC01^gHL^ B cells a cell fate differentiation into early formed B_PC_. Collectively, these results reveal a new role for IL-2 cytokine acting as a key early secreted T helper factor that participates in shaping B cells maturation and induces plasma cell differentiation.

### IL-2 acts intrinsically on CD25 expressing B cells to promote their plasma cell reprogramming *in vitro*

Although previous literature has argued against a role for IL-2 acting intrinsically on mouse B cells, our results suggest that it might directly control their cell fate commitment during antigenic stimulation and cognate interaction with T helper cells. To specifically address the role of IL-2 in controlling B cell differentiation towards plasma cells, we explored *in vitro* culture systems that would allow us to assess CSR (expression of B220, Pax5, IgG1) and B_PC_ reprogramming (expression of IRF4, Blimp-1, CD138) (Kallies et al., 2004; Sanz et al., 2019; Schena et al., 2013).

To this end, naïve B cells were isolated from wild-type or VRC01^gHL^ mice and activated with different combinations of TLR agonists, anti-BCR, anti-CD40, IL-4 and/or IL-21 (two key Tfh-secreted cytokines that support B cells survival and differentiation) (Figure 3A). We compared cell survival, plasma cell formation, downregulation of B220, and CD25 expression on each tested condition to optimize plasma cell response to various doses of IL-2 (Figure S5A-S5C). Ultimately, we observed that the integration of three signals—CpG stimulation of TLR9, anti-IgM stimulation of BCR, and IL-4/IL-21 T helper cytokine exposure—was more efficient in inducing full B_PC_ reprogramming exclusively in the presence of IL-2 (Figure 3A-3E and S5D).

**Figure 3.**
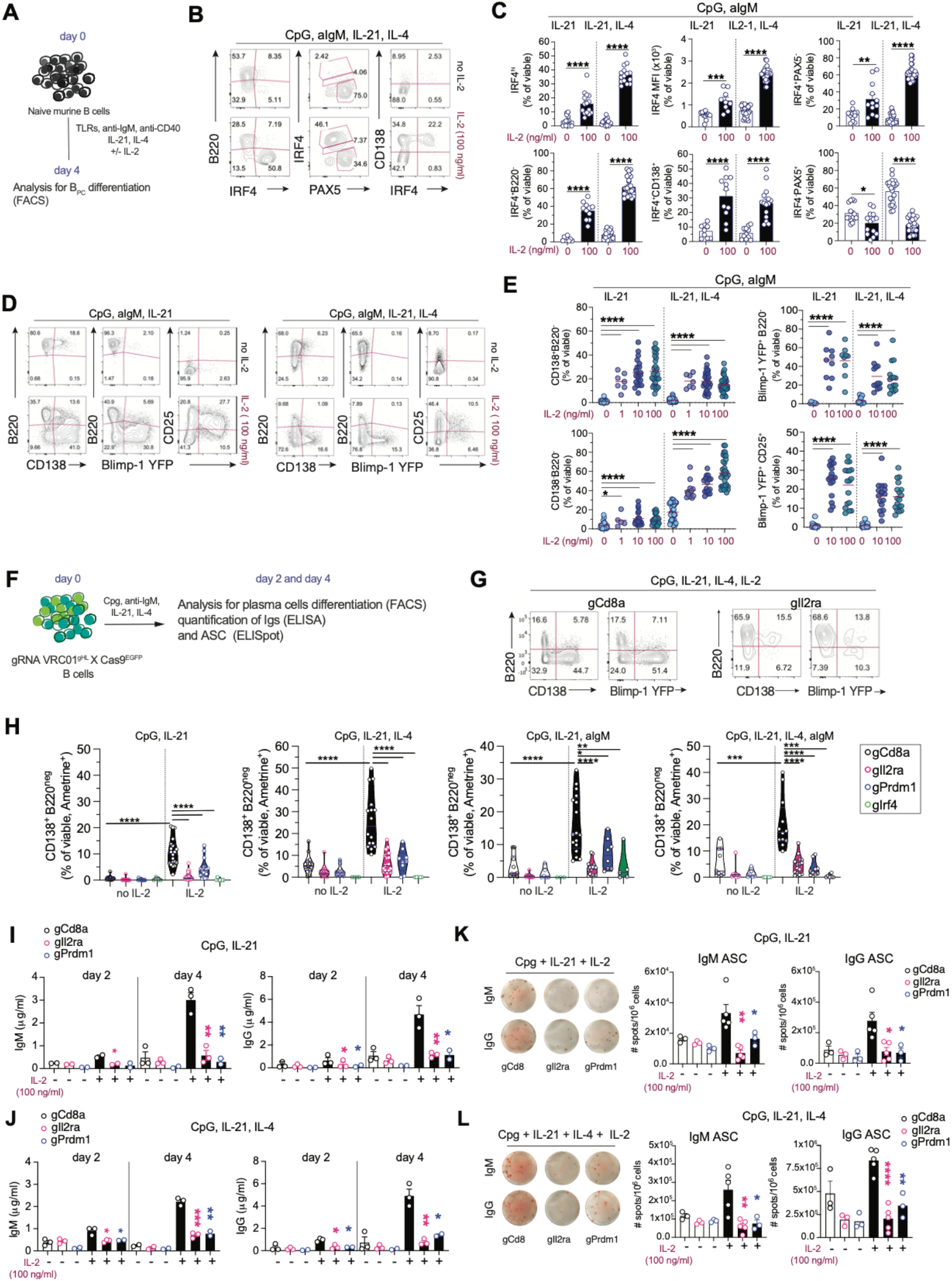
IL-2 is required to promote *in vitro* terminal plasma cell differentiation and acts intrinsically on CD25 expressing B cells. (A)Diagram showing the *in vitro* culture system and analysis performed with naïve B cells isolated from B6 or VRC01^gHL^-tg mice and stimulated *in vitro for 4 days*. (B) Representative FACS plots showing IRF4, PAX5, B220 and CD138 expression on day 4 for B cells differentiated in the presence or absence of IL-2. (C) Relative quantification of each indicated subsets and stimuli tested. (D) Representative FACS plots showing the expression of B220, CD138, Blimp-1 YFP (YFP-reporter mice) and CD25 on day 4 differentiated B cells. (E) Quantification of plasma cells differentiation for the populations of CD138^pos^B220^neg^, CD138^−neg^B220^neg^. blimp-1^pos^CD20^neg^ and blimp-1^pos^CD25^pos^ cells. Lines in graphs represent the mean value. Data are combined from n=13 individual experiments. (F) Diagram showing the *in vitro* culture system and analysis performed with CRISPR/Cas9 transduced B cells (gRNA VRC01^gHL^xCas9^EGFP^) tested for their capacity to differentiate into ASC *in vitro* and to investigate the intrinsic role of IL-2 on CD25 expressing B cells. (G) Representative FACS plots showing transduced B cells from gCd8 (control) and gIl2ra (CD25 KO) B cells differentiated into B220^−^CD138^+^Blimp-1 YFP^+^ plasma cells and (H) their quantification on day 2 post *in vitro* stimulation with the indicated cocktails of stimuli. Data are from n= 20 experiments. (I, J) ELISA assay for the quantification of secreted IgM and IgG titers in the culture supernatants collected on day 2 and day 4, for each indicated stimulation. Data are from n= 3 experiments. (K, L) ELISpot assay to enumerate the IgM and IgG ASC generated after 4 days of culture. Data are from n= 5 experiments. Each dot represents an individual mouse. Error bars represent the SEM. P values were determined by two-tailed Student’s t test. Not significant, ns p > 0.05; *p < 0.05; **p < 0.01; ***p < 0.001; ****p < 0.0001.

In sum, these results established that in our *in vitro* tested conditions the presence of IL-2 is a required factor that via the induction of IRF4 and Blimp-1 and the downregulation of B220 enable a full B_PC_ reprogramming.

### Establishment of a novel CRISPR/Cas9 editing method to generate knock-out B cells

To examine the role of IL-2 and its intrinsic regulation of B cells, a novel retroviral system was developed and validated to generate gene-specific knock-out B cells that lacked class switching or CD138 expression (Figure S6A-S6D). This new system facilitated high efficiency of transduction (Figure S6E) and selective and functional gene disruption (Figure S6F). To examine the function of IL-2 on responsive B cells, CD25 KO B cells were generated by targeting Il2ra, the gene encoding the inducible, high-affinity component of the IL-2 receptor (Figure S6G-S6I).

As done previously with Smarta cells, *Cd8a* was targeted and used as transduction controls (gCd8a B cells). Like stimulated naïve B cells, also gCd8a B cells required the addition of endogenous IL-2 to generate high levels of B_PC_ in culture (Figure 3F-3H). This requirement for exogenous IL-2 both confirms the lack of spontaneous differentiation in our transduction system and suggests that autocrine IL-2 signaling is insufficient for robust B_PC_ development. In the presence of IL-2 signals, gIl2ra B cells failed to generate B_PC_ (Figure 3G and 3H). Functionally, gIl2ra B cells showed reduced secretion of IgM and IgG (Figure 3I and 3J), as well as a reduced number of counted ASCs (Figure 3K and 3L) when compared to control cells and similarly to BLIMP-1 KO (gPrdm1) cells.

An alternative system, based on the delivery of RNP complexes into activated wild-type B cells to generate CD25 KO cells, yielded similar impediments in B_PC_ differentiation (Figure S6J-S6N).

### IL-2 signaling promotes the transition from PB to PC by sustaining high IRF4 and Blimp-1 expression, in a BACH2-independent manner

We next sought to investigate the precise mechanism by which IL-2 could control IRF4 and Blimp-1 expression in B cells. Previously, an incoherent regulatory network described in B cells identified IRF4 activity involved in controlling two mutually antagonist programs of gene expressions related to CSR/SHM and plasma cells formation (Sciammas et al., 2011). Graded expression of IRF4 can control the transition from B_GC_ to B_PC_ (Sciammas et al., 2006), with IRF4^hi^ B cells committed to becoming B_PC_, both *in vitro* and *in vivo* (Klein et al., 2006). While the previous work has attributed this process purely to BCR signal strength (Sciammas et al., 2011) the role of IL-2 in the process is less clear. Considering the importance of the addition of IL-2 in promoting B_PC_ reprogramming and inducing IRF4 and Blimp-1 expression (Figure 4A and 4B), we aimed to disentangle the role of BCR strength and IL-2 signaling.

**Figure 4.**
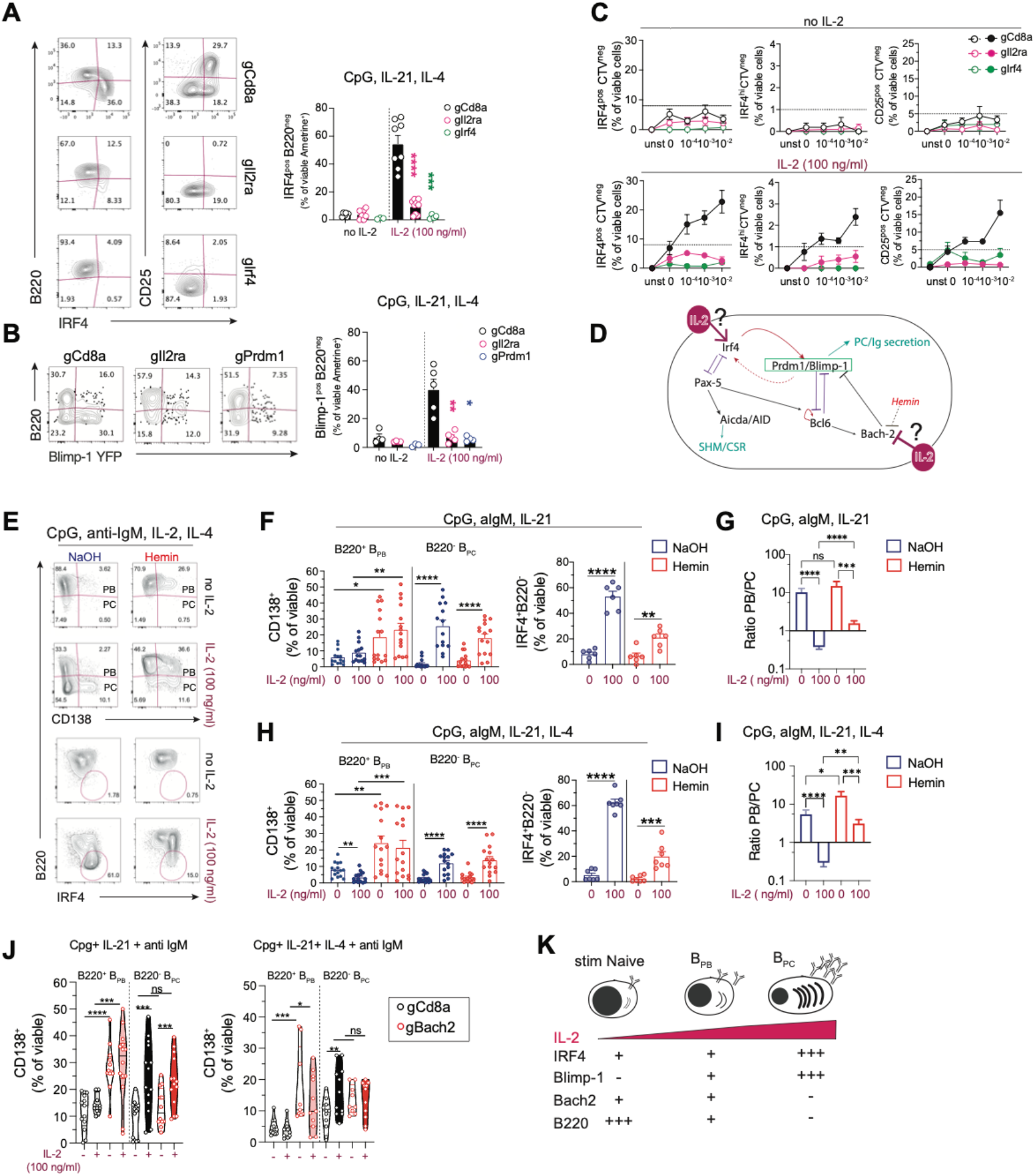
IL-2 controls the plasma cell gene regulatory network *in vitro* by inducing high expression of IRF4 and Blimp-1 in a Bach-2 independent manner. (A)FACS plots and quantification of B220^neg^IRF4^pos/hi^ PC generated upon gRNA VRC01^gHL^ stimulation. (B) FACS plots and quantification of B220^neg^blimp-1^pos^ PC generated upon gRNA VRC01^gHL^ stimulation. Data are from n= 4 or 5 experiments. (C) Test for IRF4 graded expression via BCR triggering and IL-2 signal. CTV labeled gRNA VRC01^gHL^ B cells were stimulated *in vitro* and their expression of IRF4 and CD25 was assessed on day 4. Data are from n=3 pooled independent experiments. (D) Scheme showing the two potential mechanisms involved in IL-2 regulation of IRF4 and blimp-1 and regulation of Bach-2 involvement in prdm-1 repression. (E) Dot plots showing the formation of PB and PC from stimulated naive B cells treated with Hemin (inhibitor of Bach-2) or its vehicle (NaOH) 24 hours after initial stimulation with the indicated cocktails and analyzed on day 4. (F-G) Relative quantification of PB (B220^+^CD138^+^) and PC (B220^−^CD138^+^) and IRF4 expressing cells in each indicated condition. (H, I) Ratio of PB/PC calculated from data shown in (F) and (G). Lines in graphs represent the mean value. Data are from n=4 experiments. (J) Quantification of CD138 expressing PB and PC generated upon stimulation of control gCd8 and BACH-2 KO (gBACH-2) VRC01^gHL^ cells after 2 days of stimulation with the indicated conditions. Data are from n=10 independent experiments. (K) A simple model of IL-2 graded exposure and regulation of TFs and B220 expression on stimulated naïve, PB and PC. P values were determined by a two-tailed Student’s t-test. Not significant, ns p > 0.05; *p < 0.05; **p < 0.01; ***p < 0.001; ****p < 0.0001.

We took advantage of our CRISPR-based genome editing and sought to investigate the involvement of IL-2 signaling in inducing high expression of IRF4 upon B cells activation. To this end, we tested VRC01^gHL^ x Cas9^EGFP^ B cells targeted with gCd8a, gIl2ra, or gIrf4 upon *in vitro* stimulation with anti-CD40 and IL-4. eOD-GT5-60mer was used to crosslink the BCRs and was titrated and tested with or without the addition of IL-2 (Figure S7A). CTV^lo^ proliferated B cells were analyzed on day 4 for their expression of IRF4, CD25, CD138 and Blimp-1 (Figure S7B-S7D). gIrf4 B cells were used as a control for IRF4 regulation and expression.

We observed that despite graded increase of BCR stimulation, the gCd8a control cells stimulated in the absence of IL-2 failed to express high levels of IRF4 and CD25 (Figure 4C and S7B) as well as CD138 or Blimp-1 (Figure S7C and S7D), consistent with the absence of B_PC_ formation. In support of a mandatorily required presence of IL-2 to promote B_PC_ initiation and maintenance, gIl2ra VRC01^gHL^ B cells failed to upregulate IRF4, CD138 or Blimp-1 in any tested condition and despite receiving increased BCR strengths (Figure 4C and S7B-S7D). These findings strongly suggest that increased BCR strength alone is not sufficient to promote full B_PC_ differentiation and further confirm the crucial role of IL-2 in inducing this terminal process.

As a master gene regulator of plasma cell differentiation, Blimp-1 (*PRDM1*) can be repressed by BACH2 during B cells activation (Ochiai et al., 2006). IL-2 mediated repression of BACH2 has been proposed also during *in vitro* human B_PC_ differentiation (Hipp et al., 2017), although the distinction between this operating mechanism of regulation in CD20^+^ PB versus CD20^−^ PC is unclear.

We sought to investigate the role of IL-2 in controlling the regulatory network of genes involved in SHM/CSR versus PC formation (Figure 4D). Specifically, if IL-2 acts on Bach2 as repressor-of-repressor (indirect Blimp-1 induction via IL-2), interfering with Bach2 repression should give rise to PC irrespectively of IL-2. Conversely, if the lack of Bach2 repression is not sufficient to promote fully plasma cells formation, IL-2 signal would necessitate to act upstream at the time of early irf4 induction and self-propagation of positive feedback loop of Irf4-prdm1 circuit (Figure 4D), To address this question, we tested Hemin, a potent inhibitor of BACH2 *in vitro* (Watanabe-Matsui et al., 2011). The addition of Hemin to the cultured B cells (added 24 hours from stimulation) had a minor, significant impact on cell survival when in the absence of IL-4 (Figure S8A), and as expected promoted a general increase of Blimp-1 expression in the treated B cells and irrespectively of IL-2 (Figure S8B).

The formation of B_PC_ was assessed on day 4. Here, we observed two important phenomena relative to Hemin-dependent BACH2 repression *in vitro*: first, although Hemin treatment alone was sufficient to induce the expression of IRF4 and CD138, in the absence of IL-2 it failed to promote the downregulation of B220, a hallmark of final plasma cells differentiation (Kallies et al., 2004) (Figure 4E and 4H and S8C). Hemin-treated B cells instead expressed markers of B_PB_, which represent an intermediate state of differentiation when B cells are progressing towards their final B_PC_ stage. Furthermore, while the addition of IL-2 to Hemin-treated B cells promoted the generation of some B_PC_, we observed that the ratio of PB/PC was greater than IL-2 exposed control cells (Figure 4G-4I). These findings support a mechanism by which solely inhibiting BACH2 in activated B cells without providing IL-2 generates an intermediated state of B220^+^ B_PB_ which counteracts the formation of B_PC_.

A second confirmation of IL-2 and Bach2 regulation of B_PB_/ B_PC_ transitioning was observed with gBach2 B cells (BACH2 KO). Despite increased Blimp-1 expression by Bach2-disrupted cells (Figure S8D), these cells failed to induce greater B_PC_ in the absence of IL-2 (Figure 4J) within respect to control cells. In summary, this IL-2-dependent plasma cell reprogramming relies on early induction and sustained higher expression of IRF4, which in turn regulates Blimp-1, in a BACH2-independent manner (Figure 4K).

### B cell metabolic fitness is controlled by IL-2 through the regulation of the mTOR/AKT/Blimp-1 axis

Due to established links between plasma cell reprogramming and mTOR activity (Boothby & Rickert, 2017), we hypothesized that IL-2-dependent signals might control the expression of Blimp-1 in B cells via mTOR regulation (Figure 5A), as previously reported for CD4 T cells (Ray et al., 2015).

**Figure 5.**
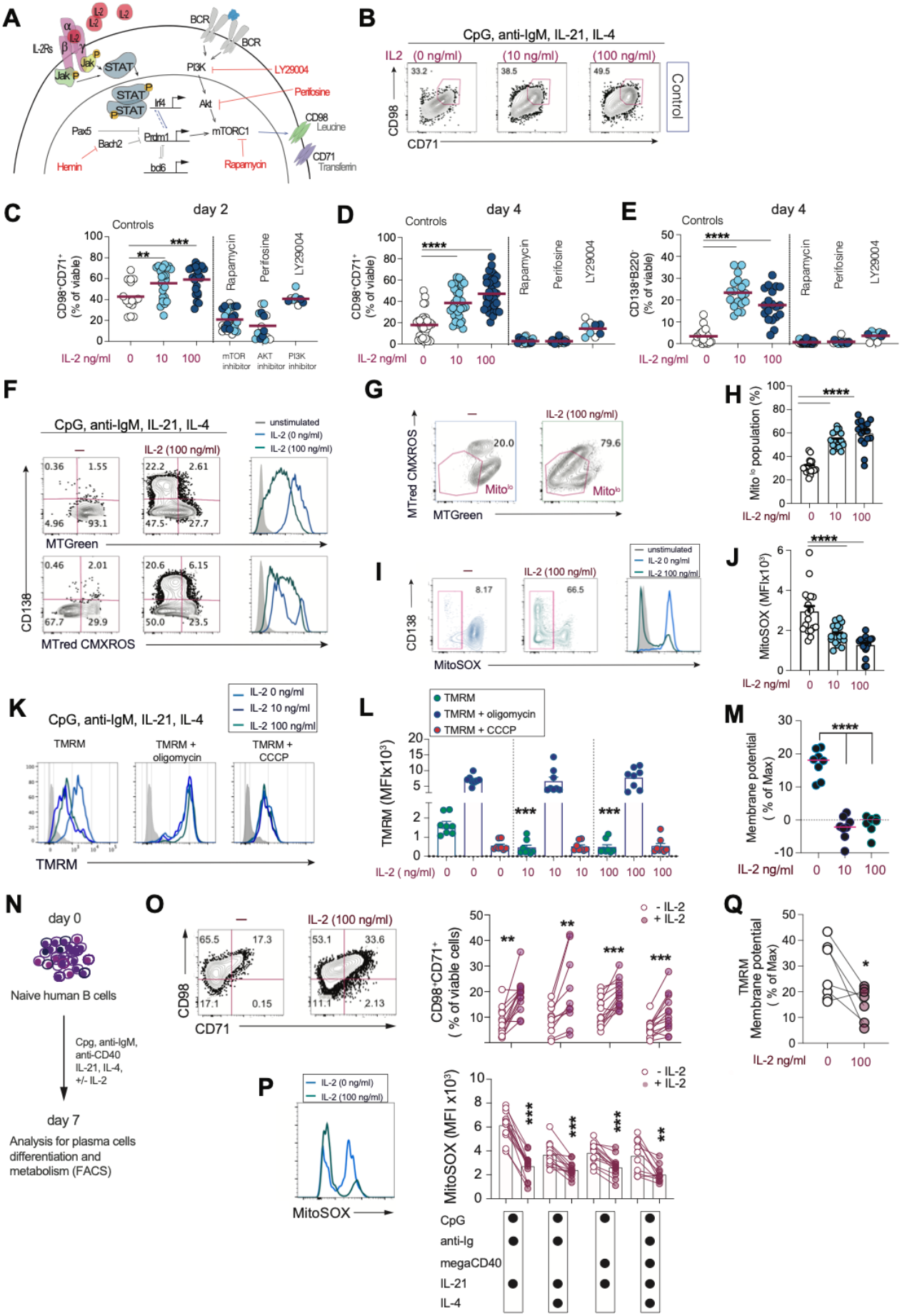
IL-2 acts via the mTOR/AKT/Blimp-1 axis to control the B cell metabolic state. (A) Proposed model of lL-2/IL-2Rs signaling and regulation of TFs, mTOR/AKT/Blimp-1 axis and CD98, CD71 expression upon B cells stimulation. Inhibitors drugs used in the experiments are shown in the model. (B) Representative dot plots showing the expression of CD98 and CD71 on day 4 post-stimulation in stimulated B cells without inhibitors (control). Increasing doses of rIL-2 (0, 10 and 100 ng/ml) are shown. (C) Quantification of CD98^+^CD71^+^ B cells on day 2 and day 4 (D) post-stimulation of B cells “controls” or treated with the indicated inhibitors. (E) Analysis of plasma cells differentiation on day 4 for each condition indicated. Data are shown for all the tested inhibitors targeting PI3K (LY294002, DMSO control), AKT (Perifosine, H^2^O control) and mTOR (Rapamycin, DMSO control), in comparison to vehicle-treated controls. Data from each control were pooled together since the results were similar. Data from inhibitors ad IL-2 (0, 10 and 100 ng/ml) were pooled since the results were similar. Data are from n= 5 experiments. Error bars represent the SEM. (F) FACS plots showing the expression of MitoTracker Green (MTG), MitoTracker Red CMXRos (CMXRos), and their MFIs compared to unstained control (day 4 of culture). (G) Representative dot plots and (H) quantification of Mito^low^ cells on day 4. (I) Representative dot plots and (J) MFI quantification of MitoSOX expressing cells analyzed after 4 days. Data are from n=5 individual experiments. Error bars represent SEM. (K) FACS plots for TMRM staining (alone, with oligomycin or CCCP) to measure membrane potentials in stimulated B cells. (L) TMRM values for each condition. (M) Percent of the maximum membrane potential used by B cells, calculated using the algorithm: [100 × (MFI ^(TMRM alone)^ - MFI ^(TMRM+FCCP)^)/(MFI ^(TMRM + Oligomycin)^ - MFI ^(TMRM+FCCP)^)]. Data are pooled from n= 3 experiments. (N) Scheme of human B cells differentiation into plasma cells and analysis. (O) FACS plots showing the expression of CD98 and CD71 on B cells after 7 days of culture and relative quantification of CD98^+^CD71^+^ cells for each indicated combination of stimuli. (P) Representative plot for the MFI of MitoSOX of B cells stimulated without or with IL-2 and relative quantification for each indicated combination of stimuli. (P) Percent of the maximum membrane potential used by human B cells. Data are pooled from n= 3 experiments pooling results from naïve B cells isolated from 13 HDs samples. P values were determined by paired Wilcoxon test. Not significant, ns p > 0.05; *p < 0.05; **p < 0.01; ***p < 0.001; ****p < 0.0001.

Consistent with our hypothesis, the addition of IL-2 to *in vitro* cultures of stimulated naïve B cells increased the frequency of cells expressing phosphoS6^(S235/236)^ and phosphoAKT^(S473)^ (Figure S9A and S9B) as well as nutrient transporters CD98 and CD71 (Figure 5B and 5C), within 48 hours at levels sustained until the B_PC_ were fully formed on day 4 (Figure 5D and 5E, and S9C and 9D).

Conversely, B cells stimulated in the presence of selective inhibitors of the PI3K/AKT/mTOR pathway (PI3K - LY29004; AKT - Perifosine; mTOR - Rapamycin) showed variable and predictable effects on B cells survival (Figure S9E) with a significant reduction in the phosphorylation of S6 and AKT, nutrient transporter expression, and B_PC_ formation visible as early as 2 days post stimulation (Figure 5C-5E, and S9A and S9C). Of note, we observed that on day 4 the B cells that co-expressed CD98 and CD71 were also enriched in the CD25 and Blimp-1 positive B cells (Figure S9F-9G).

Treatment of B cells with UK5099, an inhibitor of mitochondrial pyruvate import involved in LLPC maintenance *in vivo* (Lam et al., 2016), did not affect B cell survival (Figure S9H) or frequency of CD98^+^CD71^+^ B cells, but still showed reduced B_PC_ formation (Figure S9I). This result would suggest that while the IL-2 mediated B_PC_ formation is partially dependent on pyruvate import into the cells, inhibiting this transport does not seem to affect the mTOR induction of nutrient transporters in our short-term *in vitro* culture system.

Mitochondrial functions are associated with B cell fate and differentiation, with B_PC_ state characterized by low mitochondrial mass and potential and reduced ROS accumulation (Jang et al., 2015). To examine the metabolic profile of IL-2 responsive B cells, we implemented metabolic-stress tests. We specifically evaluated total mitochondrial mass and membrane potential using MitoTracker Green and MitoTracker red CMXROS dyes, respectively. Additionally, TMRM staining - in the presence or absence of oligomycin and CCCP - was tested to further investigate the membrane potential of activated B cells. Four days after stimulation, IL-2 treated B cells were mainly comprised of a Mito^lo^ population with reduced ROS accumulation and membrane potential (Figure 5F-5M). B cells unexposed to IL-2 stained high for MitoTracker dye, accumulated high levels of ROS and membrane potential, all features consistent with lack of B_PC_. However, this absence of B_PC_ did not appear to be dependent on the excessive ROS accumulation, since treatment of these cells with MitoTEMPO (a mitochondrial targeting compound with antioxidant properties), albeit reducing ROS accumulation could not rescue the formation of B_PC_ (Figure S9J).

We next wondered whether IL-2 could control human B cell metabolic fitness similarly to murine B cells. While the role of IL-2 in inducing B_PC_ in human B cells is well established, no direct connection has been made on IL-2 mTOR regulation. To test our hypothesis, we repeated the conditions previously tested on mouse cells, and combined different stimuli - CpG, anti-Ig, anti-CD40, IL-21, and IL-4 - to differentiate *in vitro* human naïve B cells isolated from healthy donors (Figure 5N). As expected, within a week post-stimulation, IL-2 signals promoted the formation of CD20^lo/neg^ IRF4^+^CD27^+^CD38^+^ B_PC_ (Figure S10A-10C). Notably, as observed with activated mouse B cells, also stimulated human B cells increased their expression of CD98 and CD71 in response to IL-2 (Figure 5O and S9D and S9E). This enhanced B_PC_ formation and increased mTOR activation, as observed with mouse B cells, was accompanied by a significant reduction of both ROS accumulation (Figure 5P) and mitochondrial membrane potential (Figure 5Q).

Altogether, these results show for the first time that IL-2 signaling can instruct and dictate the metabolic state of differentiating B cells via early mTORC1 regulation and induction of plasma cell differentiation. Collectively, we discovered a mechanism in which IL-2 promotes B cell metabolism and induces Blimp-1 through the regulation of mTOR activity and mitochondrial remodeling of activated B cells, which appeared to be conserved across species.

### IL-2 secreting Smarta T helper cells control the early induction of EF VRC01^gHL^ B_PC_

We next asked whether the role of IL-2 in promoting *in vitro* PC differentiation was relevant to *in vivo* processes where antigen reactive B cells could be instructed by several signals to differentiate towards EF or GC-oriented trajectories in a T cells dependent manner. Our CRISPR/Cas9 transduction system allowed us to follow the maturation of transduced VRC01^gHL^ x Cas9^EGFP^ B cells *in vivo* during early differentiation and late memory responses (Figure S11A-S11F). We validated our *in vivo* transfer system with transduced B cells by targeting genes with known expected outcomes in B cells, such as Foxo1 (gFoxo1), Hvem (gHvem) or Blimp-1 (gPrdm1). *In vivo* tested KO B cells led to the expected results. FOXO-1 KO VRC01^gHL^ B cells had an impaired DZ localization (Dominguez-Sola et al., 2015) (Figure S11G and S11H). HVEM KO VRC01^gHL^ B cells showed increased GC B cells competitiveness (Mintz et al., 2019) when transferred in competition with gCd8a control cells (Figure S11I and S11J). Blimp-1 KO VRC01^gHL^ B cells were characterized by reduced B_PC_ formation, (Figure S11K and S11L) and antibody secretion (Figure 6I and 6J) but increased GL7 expressing IgG1^+^ cells (Figure 6G and 6H and S11N) (Shapiro-Shelef et al., 2003).

**Figure 6.**
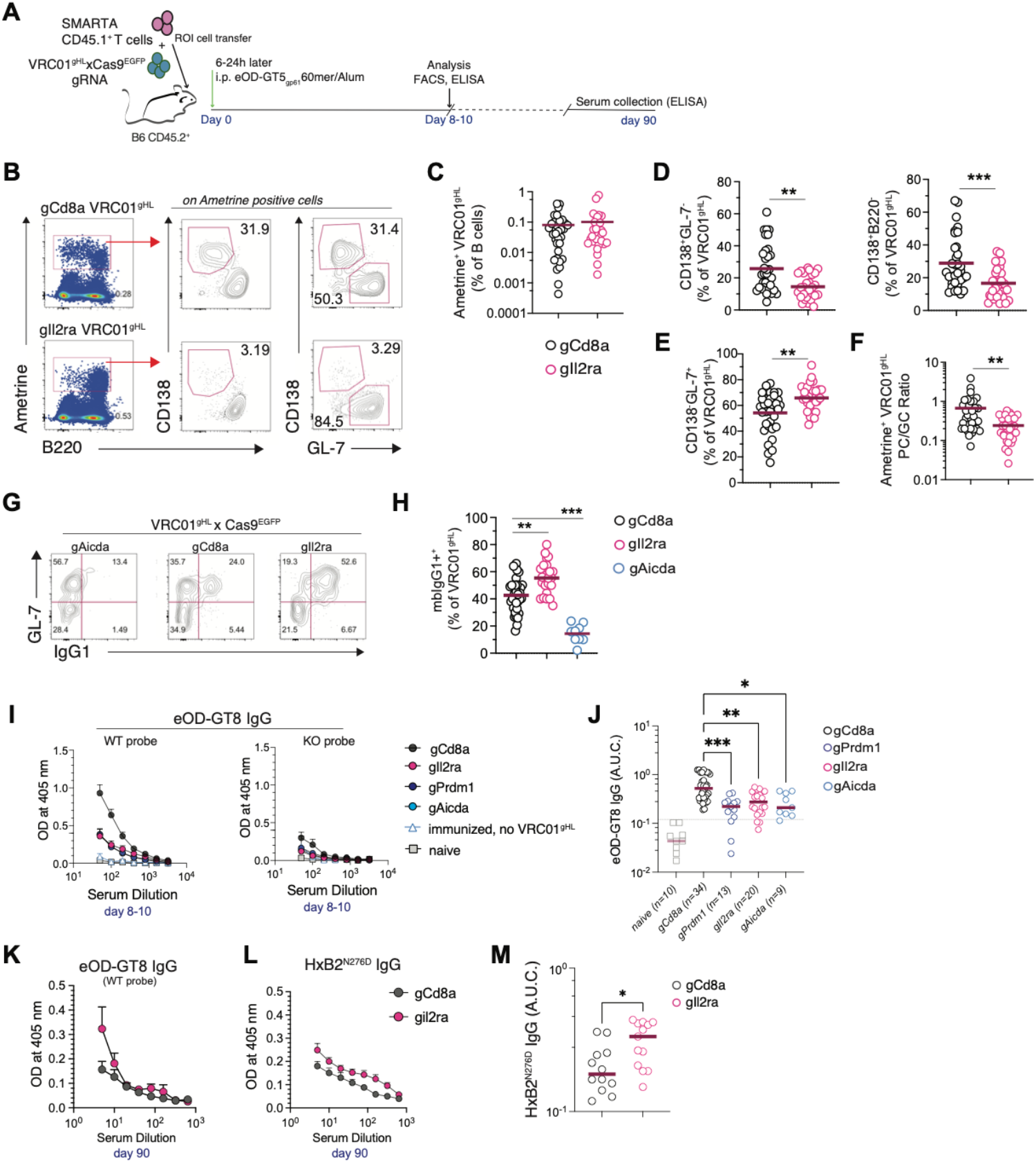
*In vivo* IL-2 signals control EF BPC versus GC differentiation and affinity maturation of antigen-specific IgG. (A) Schematic representation of adoptive cell transfer of Smarta T cells CD45.1^+^ (50K cells transferred/mouse) and transduced VRC01^gHL^xCas9^EGFP^ B cells (0.5-2×10^6^/mouse) into wild type B6 CD45.2^+^ recipient mice. Mice were immunized with 20 µg of eODGT5^gp61^60mer in Alum and spleens analyzed at 8-10 days (early plasma cells formation). Sera were tested in ELISA for antibody maturation at a memory time point (3 months post-immunization). (B) FACS plots showing the gating strategy to quantify VRC01^gHL^xCas9^EGFP^ transduced B cells (Cas9^+^Ametrine^+^) and their differentiation into B_PC_ (B220^−^GL-7^−^CD138^+^) or B_GC_ (GL-7^+^CD138^−^). (C). Quantification of gCd8 and gIl2ra VRC01^gHL^xCas9^EGFP^ frequency in spleens scored on day 7 (D) Quantification of PC VRC01^gHL^ cells as CD138^+^GL7^−^ and CD138^+^B220^−^ cells. (E) Quantification of GC VRC01^gHL^ cells as CD138^−^GL7^+^. (F) Ratio of PC/GC of Ametrine positive VRC01^gHL^ transferred cells. (G) FACS plots showing the quantification of IgG1 positive cells. (H) Analysis of membrane bound IgG1 (mbIgG1) VRC01^gHL^ B cells from gCd8 and gIl2ra B cells. gAicda VRC01^gHL^xCas9^EGFP^ are shown as reference controls for the targeted reduced CSR to IgG1. Data are combined from n= 5 independent experiments. Line and error bars represent the mean. Each dot represents an individual mouse. (I) ELISA assay to test circulating reactive IgG with eOD-GT8 and (J) A.U.C. quantification for the indicated mice. (K) GT8 and HxB2^N276D^ IgG reactive from sera of immunized mice collected on day 90 post-immunization. Data on day 90 are from mice recipients of sorted (Ametrine^+^) gRNA VRC01^gHL^xCas9^EGFP^ transduced B cells. Each dot represents an individual mouse. P values were determined by two-tailed Student’s t test. Not significant, ns p > 0.05; *p < 0.05; **p < 0.01; ***p < 0.001; ****p < 0.0001.

To directly investigate the differentiation of CD25 KO B cells *in vivo*, we adoptively transferred retrovirally transduced gCd8a or gIl2ra VRC01^gHL^ x Cas9^EGFP^ B cells with Smarta T cells into wild-type B6 recipient mice. The host mice were then immunized with eOD-GT5_gp61_60mer immunogen in alum (Figure 6A). In this set of experiments, gCd8a VRC01^gHL^ x Cas9^EGFP^ B cells were used as controls. Mice receiving gPrdm1 and gAicda (*Aicda*, AID) VRC01^gHL^ x Cas9^EGFP^ B cells were used as targeted reference genes for reduced B^PC^ formation and CSR, respectively. At 8-10 days post-immunization, AID KO VRC01^gHL^ and gPrdm1 VRC01^gHL^ B cells had significantly reduced numbers of B_PC_, and no significant impairment in the number of GC-like VRC01^gHL^ B cells (Figure S11 M).

For CD25 KO (gIl2ra) VRC01^gHL^ x Cas9^EGFP^ B cells, assessment of differentiation into either B_GC_ or B_PC_ cells (Figure 6B) revealed that a smaller frequency of cells were capable of differentiating into B_PC_ (Figure 6C-F), though absolute numbers of VRC01 B_PC_ were unchanged (Figure S11 M). CD25 KO cells preferentially differentiated along the GC pathway and were poised to class-switching into IgG1 expressing B cells (Figure 6G and 6H), similarly to BLIMP-1 KO VRC01 B cells.

Functionally, the mice transferred with CD25 KO VRC01^gHL^ B cells showed a significantly reduced accumulation of eOD-GT8 IgG, similarly to BLIMP-1 and AID KO cells (Figure 6I-6K). After 3 months, mice receiving CD25 KO cells had some detectable IgG to eOD-GT8, and possibly HxB2^N276D^ (Figure 6K-6M). Overall, the data support the conclusion that early exposure of activated antigen-specific B cells to IL-2 provided by cognate T helper cells preferentially instruct B cells to mature as early EF B_PC_.

## DISCUSSION

A direct role for IL-2 in controlling Blimp-1 expression and consequently modulating B cell metabolic fitness has not been fully investigated. The current literature is discordant in reference to an intrinsic role for IL-2 in the regulation of B cells. Studies with Il2ra/CD25 knock-out B cells have been limited by abnormal lymphocyte maturation and autoimmune phenotype of Il2ra^−/-^ mice (Willerford et al., 1995). Despite early reports that IL-2 could be inducing Blimp-1 expression *in vitro* (Turner et al., 1994), follow-up studies performed in mice were mainly focused on the understanding of the role of IL-2 signaling in the regulation of Tfh and Tfr cells, and eventually discounted a direct role for IL-2 on B cells (Ballesteros-Tato et al., 2012; Botta et al., 2017).

In our study, we took advantage of CRISPR/Cas9 genome-based editing (Ran et al., 2013) and established an efficient protocol of transduction for T and B cell-specific gene disruption. By complementary targeting of the IL-2/IL-2R pair on IL-2 secreting Smarta cells or IL-2R expressing VRC01^gHL^ B cells, we instituted a system that allowed us to perform both *in vitro* and *in vivo* studies and enable precise investigation of IL-2 signals on antigen-specific immune cells in the context of immunization. This system can be used in the future to investigate the function of specific molecules and paired-couple targets expressed by both T and B cells. Additionally, this model of immunization with HIV-related immunogens and adoptive transfer of antigen reactive Smarta T and VRC01^gHL^ B cells is relevant for the understanding of T cell help on B cells that can be translatable to vaccine-mediated responses in current clinical trials with eOD-GT8 HIV candidate in humans (Venkatesan, 2021).

Overall, our results provide clear pieces of evidence of intrinsic regulation of IL-2 on CD25 expressing B cells. We uncovered a mechanism dependent on IL-2 signaling able to induce the two master transcription factors Irf4 and Blimp-1 in activated B cells. We found that Blimp-1 activity, while promoting plasma cell reprogramming, was not influenced by BACH2 repression but required direct signals from IL-2. Additionally, we observed that IL-2 direct induction of Blimp-1 facilitated metabolic reprogramming of B cells through an mTORC1-dependent regulation of the energetic machinery of fate-committed B_PC_. A similar mechanism was observed with cultured human B cells, suggesting that IL-2 can regulate different human B cell subsets with respect to their mTORs-related sensitivity and Blimp-1 dependency.

In our working model, we hypothesized that the mechanism by which IL-2 induces Blimp-1 might occur in two ways: (1) IL-2 indirectly induced Blimp-1 via BACH2 or (2) IL-2 directly induced Blimp-1 in a mechanism that is not contingent on BACH2 mechanisms of repression. Since the induction of B_PC_ in the absence of IL-2 could not be achieved by solely repressing BACH2, the mechanism proposed in (1) does not appear to be occurring and should be excluded in favor of (2). In fact, the data strongly suggest against a co-dependency of IL-2 and BACH2 for the regulation of Blimp-1, since the combined addition of IL-2 and Hemin to the culture did not synergically increase the frequency of B_PC_ but instead appear to counteract the progression of B_PC_ over B_PB_.

Our data support a model where early exposure to IL-2 during cognate T-B cell interactions favors the decision of activated B cells to commit into antibody-secreting B_PC_ and progress their differentiation at EF sites. In *vivo*, the induction of IL-2 and its receptor occurs early upon immunization (Fuhrmann et al., 2016). Therefore, we speculate that upon antigen encounter, early formed IL-2 secreting T helper cells (pre-T_FH_) capable of initiating crosstalk with B cells at the T-B border instruct IL-2 responsive B cells to mature away from the trajectory of GC and SHM processes. These B cells will undergo instead drastic metabolic changes and initiate their fate reprogramming through the upregulation of IRF4 and Blimp-1, and the activation of mTOR in support of terminal plasma cell differentiation.

IL-2 functions are well established in the context of infections or vaccinations as well as during aberrant autoimmune disease manifestations, conditions where B cells can either execute central protective functions or undergo break-of-tolerance (Ahmed et al., 2019) with concomitant aberrant T helper cells responsible of immunopathology (Faliti et al., 2019.; Weinstein et al., 2012). For instance, IL-2 secretion has been reported in the case of acute infection with SARS-2 (Costela-Ruiz et al., 2020), a viral infection where exaggerated EF responses have been correlated with morbidity (Woodruff et al., 2020), similarly to the aberrant effector functions of EF B cells that can drive immunopathology in systemic lupus disease (Lieberman & Tsokos, 2010; Woodruff et al., 2020).

Due to its potent immunomodulatory functions, IL-2 has been also evaluated for clinical purposes in disparate settings, and in a recent study (Yshii et al., 2022) Yshii and colleagues elegantly described a brain-specific IL-2 delivery system capable of protecting mice from pathological neuroinflammation of the central nervous system, a non-lymphoid tissue where also B cells can be recruited and play critical roles (Ahn et al., 2021).

In view of the important roles of IL-2 in shaping T and B cell responses, supporting proliferation and regulating cell fates, cell-specific targets and modulation of IL-2 signaling for therapeutical purposes should be carefully considered (Ahn et al., 2021; Ballesteros-Tato & Papillion, 2019; Papillion & Ballesteros-Tato, 2021).

In summary, our study established a new role for IL-2 in promoting the reprogramming of B cells into antibody-secreting cells during *in vivo* immunization and does have important implications for B cells biology understanding and vaccine design. Additionally, our understanding of IL-2 intrinsic regulation of B cell metabolism and functions paves the way to unexplored areas of investigation where the cytokine milieu can shape the humoral B cells responses both in mouse models and human-related B cell biology.

## ACKNOWLEDGMENTS

We would like to thank members of the LJI FACS, and DLAC Core Facilities for their outstanding expertise. We thank the LJI Clinical Core, for healthy donor enrollment and blood sample procurement and all the volunteers that participated in the enrollment studies. We thank Dr. J.H. Lee for fruitful discussions and advice on VRC01-related assays. We would like to thank Dr. M.C. Woodruff for the critical reading of the manuscript and helpful discussions.

This work was funded by the NIH P01 AI145815, CHAVD-ID UM1AI100663, and CHAVD UM1AI144462.

## AUTHOR CONTRIBUTIONS

C.E.F. established the B cells CRISPR/9 retroviral transduction and RNP electroporation protocols used in this study, performed most *in vitro* and *in vivo* experiments, flow cytometry, CRISPR/9 experiments, ELISA and ELISPOT, and data analysis. M.M. provided technical assistance and performed *in vitro* experiments, retroviral transductions, flow cytometry, ELISA, and ELISpot and generated data supervised by C.E.F. J.Y.C. and S.B. provided technical advice and expertise for molecular biology protocols. W.R.S provided reagents. C.E.F. wrote and edited the manuscript. S.C. edited and wrote the manuscript. C.E.F and S.C. conceptualized the project. S.C. provided research supervision.

## RESOURCE AVAILABILITY

### Lead Contact

Further information and requests for resources and reagents should be directed to and will be fulfilled by the Lead Contact, Shane Crotty.

### Materials Availability

The reagents generated in this study may be made available on request upon completing a Materials Transfer Agreement.

### Data and Code Availability

FCS files and imaging data generated in the current study are available from the corresponding author on request. gRNA design and sequences are listed in Table S1.

## EXPERIMENTAL MODEL AND SUBJECT DETAILS

### Mice

Animal care and mouse work were performed at La Jolla Institute for Immunology (LJI) following the guidelines of the IACUC committee of LJI. Experiments were carried out using sex- and age-matched mice of ~8-12 weeks of age. C57BL/6J (JAX: 000664) and B6.SJL-*Ptprc*^*a*^*Pepc*^*b*^/BoyJ (B6.CD45.1) (JAX: 002014) mice were purchased from Jackson Laboratory at 6 weeks of age and housed at LJI. Mouse strains described below were bred and housed in specific-pathogen-free conditions. VRC01^gHL^ mice carrying inferred germline reverted VRC01 IgH (VRC01^gH^) and VRC01 IgL (VRC01^gL^) (Abbott et al., 2018) were maintained on a C57BL/6J or B6.CD45.1 background and maintained as either homozygous or heterozygous lines. Additional lines were generated in-house by crossing homozygous VRC01^gHL^ mice to Rosa26^Cas9^ knock-in B6 mice (JAX: 028555) to generate VRC01^gHL^ x Cas9^EGFP^ Tg mice; homozygous VRC01^gHL^ mice were also crossed to Blimp-1 YFP mice (bred in-house) to generated VRC01^gHL^ x Blimp-1 YFP mice. SMARTA mice (TCR transgenic for I-A^b^-restricted LCMV glycoprotein 66–77 peptide (gp_66_)) were maintained on a B6 or B6.CD45.1 background and crossed to Rosa26^Cas9^ knock-in B6 mice (JAX: 028555,) to generate Smarta x Cas9^EGFP^ Tg mice.

### Human samples

Blood collection from healthy volunteers was approved by the Institutional Review Board of La Jolla Institute for Immunology (LJI; Normal Blood Donor Program). Whole blood was collected via phlebotomy in acid citrate dextrose (ACD) serum separator tubes (SST) or ethylenediaminetetraacetic acid (EDTA) tubes and processed for peripheral blood mononuclear cells (PBMCs), serum, and plasma isolation. Peripheral blood mononuclear cells (PBMCs) were isolated from blood by density-gradient centrifugation using Ficoll-Paque Plus gradient (GE Healthcare). Isolated PBMCs were cryopreserved in 10% DMSO freezing media and stored at −80C until *in vitro* analysis.

## METHOD DETAILS

### Flow Cytometry

Single-cell suspensions of lymphocytes were generated by standard gentle mechanical disrupting of the mouse spleens in R10 complete medium (RPMI 1640 (Corning, Catalog #10-041-CV) + 10% FBS Hyclone (GE Healthcare, Catalog # SH30071), supplemented with 2 mM GlutaMAX (Gibco, Catalog #35050-061), 50 μM β-mercaptoethanol (2-βME), 100 U/ml penicillin/streptomycin (Gibco, Catalog #15140-122), sodium pyruvate (Corning, Catalog #25-000-CI), and non-essential amino acids (Gibco, Catalog #11140-050). Isolation of cells from bone marrow was obtained by standard gentle flushing of cells from mouse femur and tibia, after careful removal of muscle and tissues. Bones were cut at both ends with sharp sterile scissors and cells were flushed with a 23G needle and ~5-8 ml of RMPI 10% FCS. Cells were filtered through a 70 μm nylon cell strainer and collected for further analysis.

Cell suspensions were centrifuged at 550 g, 4°C, for 5 minutes and pellets were resuspended in Ammonium-Chloride-Potassium (ACK) lysing buffer (Thermo Fisher Scientific, Catalog # A1049201) to lyse RBC (erythrocytes removal performed at R.T. for 5 minutes). ACK solution was blocked with an equal volume of R10 and centrifuged at 550 g, 4C, for 5 minutes. Pellets were resuspended in R10, cells counted under the microscope and stained in FACS buffer (PBS, 2% FCS, 2mM EDTA) with purified Rat Anti-Mouse CD16/CD32 (Mouse BD Fc Block™, BD Catalog #553141) and the specific antibody cocktail. Cells were stained in U-bottom 96-well plates at room temperature for 45 minutes. After the extracellular staining, dead cells were stained using Fixable Viability Dye eFluor-780 (Thermo Fisher Scientific, Catalog # 65-0865-18) in PBS. Cells were washed, resuspended in FACS buffer and acquired with flow cytometry.

Biotinylated antibodies were detected using fluorescently labeled streptavidin. Fluorescent antigen probes were used to detect VRC01^gHL^ B cells with eOD-GT8 60-mer conjugated to Alexa Fluor 647 (Thermo Fisher Scientific, Catalog # A20173). Intracellular staining was performed with a fixation kit (BD Bioscience, Catalog # 554714) following the manufacturer’s instruction.

Stained samples were acquired on a FACS Celesta (BD Bioscience) or LSR Fortessa (BD Bioscience) running FACS Diva (BD Bioscience). FCS data were analyzed using FlowJo 10 (FlowJo).

### Adoptive cell transfer, immunization with eOD-GT5_gp-61_ 60mer and in vivo treatment with IL-2

For adoptive cells transfer of Smarta cells, 0.5 × 10^5^ naïve CD4 T cells isolated by negative selection were transferred intravenously into recipient mice. Naïve B cells were isolated by negative selection. In the experiments of co-transfer with naïve CD4 T cells, VRC01^gHL^ uGFP naïve B cells were isolated and co-injected into recipient mice at low (1 × 10^3^) or high (1 × 10^6^) precursor frequency numbers. Animals were immunized the next day with the immunogen eOD-GT5^gp61^ 60mer adsorbed in Alum.

Activated, retrovirally transduced CD4 T cells were acquired on FACS Celesta after their expansion *in vitro*, to check their viability and the frequency of Ametrine positive transduced cells (ranging from 75 to 90%). For the *in vivo* transfer experiments, wells from the same gRNA were combined, cells spun down and then either directly transferred into recipient mice or sorted based on Cas9-GFP and Ametrine double expression. Transferred cells were rested into recipient mice for 3 days prior to immunization. B cells were transferred intravenously 24 h prior to immunization.

Activated, retrovirally transduced VRC01^gHL^ x Cas9^EGFP^ B cells were acquired on FACS Celesta 24 h after the second step RV to check their viability and the frequency of Ametrine positive transduced cells (ranging from 45 to 90%). For the *in vivo* transfer experiments, wells from the same gRNA were combined, cells spun down, and then either directly transferred into recipient mice or sorted based on Cas9-GFP and Ametrine double expression. RV gRNA VRC01^gHL^ x Cas9^EGFP^ B cells were co-transferred with CD45.1^+^ naïve Smarta CD4 T cells and host mice were immunized after ~6-24 h post-transfer (note that this window of immunization gave similar results). Recipient mice were immunized intra-peritoneally (i.p.) with a 200 μl solution composed of 1 part of Alum (Invivogen, Alhydrogel® adjuvant 2%, Catalog #vac-alu-250) and 1 part of diluted immunogen eOD-GT5^gp61^ 60mer, used at 20 μg in PBS. When recombinant IL-2 was tested *in vivo*, immunized recipient mice were treated daily – from day 3 to day 6 post-immunization – and received 50 ul injected intravenously (ROI, retro-orbital injection) of either rec. human IL-2 (50U/ml) or vehicle only (0.1M Acetic Acid).

### *In vitro* B cell differentiation culture system

Naive murine or human B cells were isolated by negative selection (Stemcell EasySep kit) and stimulated in B cells complete medium (R10, RMPI 1640 10% FBS Hyclone complete medium) supplemented with ITS (1:100, Thermo Fisher, # 41400-045). To stimulate murine B cells, splenocytes cells from B6 wild-type mice or VRC01^gHL^ mice strains were isolated by negative selection (Stemcell, Catalog #19854) and seeded at a density of 1×10^5^ B cells in 0.5 ml final volume of 24 well plates. B cells were stimulated with different stimuli, as indicated. B cells were stimulated with different combinations of CpG ODN 1826 (Invivogen, Catalog # tlrl-1826) used at 2.5 ug/ml; LPS (Sigma, Catalog # L5418-2ML) used at 1 μg/ml; anti-IgM (Jackson Imm. AffiniPure F(ab’)_2_ Fragment Goat Anti-Mouse IgM, µ chain specific, Catalog # 115-006-075) used ad 2.5 μg/ml; anti-CD40 (Biolegend, Cat#102812) used at 10 μg/ml; rec. human IL-21 (Peprotech, Catalog #210-21) used at 20 ng/ml; rec. murine IL-4 (R&D, Catalog #204-IL); rec. human IL-2 (Peprotech, Catalog #200-02). When indicated, Hemin (Sigma, Catalog #9039) was added after 24 hours from the initial stimulation of B cells at a final concentration of 60 μM, and controls cells were stimulated with an equimolar amount of NaOH (vehicle for Hemin).

When RV B cells were used, after the 2-steps RV transduction cells were washed twice to remove the anti-CD40 antibody and seeded at a density of 0.1 × 10^5^ cells per well in 24 well plates. The same stimulation conditions used for naïve B cells were applied to this culture. To test the regulation of IRF-4 in the presence of IL-2 and increased stimulation of BCR, CTV (Thermo Fisher, Catalog # C34557, used at 5 uM) labeled RV gRNA VRC01^gHL^ Cas9^EGFP^ B cells were stimulated with increased doses of eOD-GT5_gp61_ 60 mer, in the presence of anti-CD40 and rec. murine IL-4. Cells were analyzed on day 4 for PC formation, proliferation, and upregulation of intracellular IRF-4 and membrane-bound CD25.

To stimulate human B cells, frozen vials of PBMC from healthy donors were thawed, and naïve B cells were isolated by negative selection (StemCell Technology, Catalog #19254). B cells were seeded in 24 well flat-bottom plates at a density of 1 × 10^5^ per well and stimulated in 1 ml of B cells complete medium. Combination of each stimulus tested to the culture are indicated in each figure, and listed in here: anti-Ig (M+G+A) (Jackson Imm., Catalog #109-006-064) used at 2.5 μg/ml; CpG ODN 2006 (Invivogen, Catalog # tlrl-2006) used at 2.5 μg/ml; rec. human IL-2 (Peprotech, #200-02) used at 100 ng/ml; rec. human IL-4 (R&D, Catalog #204-IL-010) and rec. human IL-21 (Peprotech, #210-21) used at 20 ng/ml; megaCD40L (Alexis, Catalog #552-110-C010) used at 10 ng/ml.

### Metabolic test assays

Stimulated B cells were treated with drug inhibitors of the mTOR pathway, as indicated in the figures. When indicated, the PI3K pan-inhibitor LY294002 (Calbiochem Millipore, Catalog #19-142. Used at 10 μM, compared to DMSO treated control cells) was added to cells on day 0, within the other stimuli. When indicated, the following inhibitors were added on day 1: the Bach-2 inhibitor Hemin (Sigma, Catalog #9039. Used at 60 μM, compared to NaOH treated control cells); the mTOR inhibitor Rapamycin (Sigma, Catalog # R8781. Used at 50 nM, compared to DMSO treated control cells); the AKT inhibitor Perifosine (Tocris, Catalog #6087. Used at 20 μM, compared to H20 treated control cells). On day 2 cells were analyzed for the expression of phosphoproteins pS6 ^S235/236^ and pAKT^S473^ by flow cytometry. Primary antibody against p-S6 (Cell Signaling, Catalog #4858) and pAKT (Cell Signaling, Catalog #4060), and secondary AlexaFluor-647 conjugated Goat-anti-Rabbit IgG antibody was used for their detection.

For staining of mitochondria, stimulated B cells were collected, adjusted to a density of 1-3 × 10^5^ cells per well, and stained with an extracellular antibody cocktail of B220 BV785, CD138 BV650, and with the fixable viable dye eF780 (Thermo Fisher, Catalog #65-0865-14) for 15 minutes. Cells were then incubated with MitoTracker dyes Green (MTG; Thermo Fisher, Catalog #M7514) and Red (MTRed CMXRos; Thermo Fisher, Catalog #M7512) used at 20 nM, in PBS at 37 degrees for 30 minutes. After incubation, the cells were washed, resuspended in FACS buffer and acquired immediately with the LSR Fortessa.

MITOSOX Red mitochondrial superoxide indicator (Invitrogen, Catalog # M36008, 5 μM) was tested on stimulated B cells to measure ROS accumulation after 4 days in culture. Stained cells were incubated for 30 minutes at 37 degrees. Cells were then stained with 1 μM SYTOX Blue Dead Cell Stain for an additional 10 min incubation at 37 degrees, washed and rapidly analyzed in flow cytometry. The ROS inhibitor MitoTempo (Sigma, Catalog #SML0737. Used at 100 μM, compared to H20 treated control cells) was added on day 0 to the *in vitro* stimulated B cells to evaluate the effect of ROS accumulation on PC formation.

TMRM (Invitrogen, Catalog #T668) staining was carried out on activated B cells stimulated for 4 days. B cells were treated with 6 μM Oligomycin or 5 μM CCCP for 10 minutes at 37 degrees, and then incubated for 30 minutes with 30 nM TMRM and 1 μM Sytox at 37 degrees. Cells were acquired with flow cytometry. The percentage of the maximum mitochondrial potential was calculated using the algorithm: (100 × (MFI (TMRM alone)-MFI (TMRM+FCCP))/(MFI (TMRM + Oligomycin) - MFI (TMRM+FCCP))) as previously described (Akkaya et al., Nat Imm 2019).

### Smarta cells *in vitro* restimulation with gp-66 peptide and polarization assays

RV gRNA Smarta Cas9Tg cells were isolated from spleens of immunized mice, adjusted to 1 × 10^6^ cells in 200 μl of R10 and stimulated in 96 well RB plates for ~5h at 37 degrees. Splenocytes were incubated in the presence of Monensin (Biolegend, Catalog # 420701) and restimulated with 2 μg/ml of gp66 peptide or with a combination of PMA and Ionomycin used at 50 and 500 ng/ml, respectively.

To test IL-2 secretion and efficiency of Il2ra gene disruption, gRNA Smarta Cas9Tg cells were polarized to Th0 (forced neutral) with 10 µg/mL of anti-IFNγ, anti-IL-4, and anti-TGFβ as previously described (Eto et al, PlosOne 2011) while subjected to the 2-step transduction protocol. After transduction, cells were restimulated with PMA/Ionomycin and tested for their IL-2 secretion *in vitro*.

### ICS to detect secreted cytokines *in vitro*

Frozen PBMC or tonsils from healthy volunteers were thawed and cells were counted and adjusted to 20 × 10^6^/ml in R10. Cytokines’ secretion from CD4 T cells was assessed by incubating 1×10^6^ cells from PBMC or tonsils in 96 well U-bottom plates in a final volume of 200 μl of R10. Antigen-specific CD4 T cells responses were assessed by incubating with MPs “mega-pools” of peptides encompassing the sequenced of common human pathogens, CMV or PT. To detect intracellular CD40L expression, cells were pre-incubated with anti-human CD40 antibody (Miltenyl Biotech, pure-functional grade, Catalog # 130-094-133) used at 0.5 μg/ml, for 15 minutes at 37 degrees. Cells were then stimulated with each MPs (used at 2 μg/ml). An equimolar amount of DMSO was added to replicated control well of each sample. Cytokines’ secretion upon polyclonal stimulation of cells was assessed with TCR triggering via SEB (used at 5 μg/ml) or a combination of PMA and Ionomycin (used at 50 and 500 ng/ml, respectively). Cells were stimulated for ~6-8h in the presence of BFA (used at 10 ug/ml), before extracellular staining, fixation with BD fixation kit and intracellular detection of CD40L and cytokines. Fixed cells were acquired with LSRII or LSR Fortessa. The data in graphs are shown as DMSO subtracted values and negative values (Limits of detection (LOD)) were set equal to 0.001 in the log10 scale graphs.

### Plasmid generation, CRISPR gRNAs design and cloning

LsgA plasmid vector was generated by a modification to the LMPd-Amt retroviral (RV) vector (published in Chen et al., Immunity 2014) with the insertion of a sgRNA backbone vector (pUC57-U6-sgRNA, Genscript). CHOP-CHOP website (source: https://chopchop.cbu.uib.no) was used to select and design single-guides (sgRNA) testing the high-ranked guide sequences with the highest on-target and off-target scores. Sequences of sgRNAs used in this study are in Table 1.

Initial screening and validation of targets were performed by testing 2-3 guides designed for each gene target. The gRNA giving the highest efficiency of deletion was chosen for subsequent studies. The sgRNA was inserted into the LsgA vector using BbsI restriction sites. Forward (F1, 5’–CACCGN^(20)^–3’) and reverse (R1, 3’–C N^(20)^CAAA −5’) oligos specific for the 20-nt selected as CRISPR targets (designed with a G-C paired based added at the 5’ of the plasmid and lacking the PAM sequence) were synthesized by Integrated DNA Technologies (IDT) and inserted into the LsgA “empty” vector. Briefly, the LsgA vector was digested with BbsI-HF^®^ (NEB Catalog #R3539) restriction enzyme overnight, in the water bath at 37 degrees. Annealed oligos were ligated into the LsgA vector, for 2 hours at room temperature; the product of ligation was then transformed in E. Coli TOP10 chemically competent cells (Thermo Fisher, Catalog #C404010) following the manufacturer’s instruction. 50-200 μl of each transformed vial was spread on LB-Agar-Ampicillin plates and incubated overnight at 37 degrees. The next day, colonies were picked up and mini-prepped. The obtained plasmids were sequenced with Retrogen.^Inc.^ to confirm the sgRNA insertion and stored at −80C until Plat-E cells transfection.

### Plat-E cell transfection and retroviral transduction of Cas9^tg^ T and B cells for sgRNA delivery and CRISPR-mediated targeted gene disruption

Virions were produced by transfection of the Plat-E cells line, as previously described (Choi et al, Nat. Imm. 2020). Briefly, Plat-E were seeded overnight at a density of 0.8 × 10^6^ cells/well. The next day the cells were transfected adding ~1.3 μg of plasmid gRNA and 0.7 μg of pCL-Eco to maximize recombinant-retrovirus titers to 2 ml of transfection media (DMEM + 10% FCS, 1% L-Glu, 1% P/S). Plat-E cell supernatants were collected 24 h and 48 h after transfection, filtered through a 0.45 μm syringe filter, and stored at 4 degrees until transduction.

For T cells transduction, Cas9-Tg naïve CD4^+^ T cells were isolated from whole splenocytes by negative selection (Stemcell EasySep kit, Catalog #19765) and resuspended in R10 complete medium and rec. human-IL-7 used at 2 ng/ml. Cells, seeded at a density of 0.5 × 10^6^ cells/ml/well, were stimulated in 24-well plates pre-coated with 8 μg/ml anti-CD3 (17A2; BioXcell) and anti-CD28 (37.51; BioXcell). In this 2-steps RV transduction system, at 24 h and 48 h after stimulation, cells were transduced by adding ~1-2 ml of RV supernatants supplemented with 50 μM 2-βME and 8 μg/ml polybrene (Millipore, Cat # TR-1003-G), followed by centrifugation for 90 min at 524*g* at 37 °C. Following each transduction, the RV-containing medium was replaced with R10 + 50 μM 2-βME + 10 ng/ml of rec-human-IL-2. After 24 h from the second step transduction, CD4^+^ T cells were transferred into six-well plates in R10 + 50 μM 2-βME + 10 ng/ml of rec. human-IL-2 and expanded for 2 days in the incubator at 37 degrees. On day 3, the culture medium was replaced with R10 + 50 μM 2-βME and 2 ng/ml of rec. human IL-7.

For B cells transduction, Cas9-Tg naïve B cells were isolated from whole splenocytes by negative selection (Stemcell EasySep kit, Catalog # 19854) and seeded in 24-well flat-bottom plates at a density of 0.5-1 × 10^6^/well, resuspended in R10 complete medium supplemented with ITS (Gibco, Catalog #41400-045) and rec-human-IL-2 (Peprotech, Catalog #200-02) used at 100 ng/ml. B cells were stimulated for 2 days with 10 ug/ml of anti-CD40 (Biolegend, Cat#102812) and 10 ng/ml of rec. human-IL21 (Peprotech, Catalog #210-21). After 48h, activated B cells were retrovirally transduced with the 2 steps-RV protocol previously described for CD4 T cells. Briefly, B cells were spun down and resuspended in ~200-250 μl of R10 stimulation media, before adding ~1-2 ml of RV medium for transduction steps. Stimulation media was prepared fresh after each step of transduction to maximize efficiency. After 24 h from the second transduction, B cells were collected, washed twice, and counted under the microscope for *in vitro* assays or *in vivo* transfer. When required, transduced cells were sorted based on Ametrine and GFP double expression (FACSAria; BD Biosciences).

### Electroporation of Cas9/Synthetic RNA Ribonucleoprotein (RNP) Complexes for CRISPR/Cas9 Genome Editing of murine T and B cells

Electroporation-based transfection was performed by delivering RNP into activated cells with the MaxCyte platform. Each crRNA and ATTO-550-conjugated *trans*-activating CRISPR RNA (tracrRNA) were purchased from Integrated DNA Technologies (IDT).

Purified *Streptococcus pyogenes* Cas9-NLS protein was purchased from QB3 Macrolab of the University of California, Berkeley. crRNA and tracrRNA were duplexed by heating at 95 °C for 5 min. RNP complexes were generated by mixing crRNA–tracrRNA duplexes (240 pmol) and Cas9-NLS protein (80 pmol) for 10 min at 24–26 °C. Naïve B cells were isolated and stimulated as previously described for RV transduction. The cells were then transfected with an RNP mixture by electroporation using MaxCyte ATX using the B cell protocol (B1) using 2-8 × 10^6^ cells per chamber. After electroporation, B cells were transferred into 24-well plates and stimulated to promote plasma cells formation, as indicated. Transfection efficiency (ATTO550 intensity) and cell viability were measured after 24 h using LSRII or LSR Fortessa. RNP transfection efficiencies were consistently greater than 90%, with high cell viability reached within independent experiments.

### ELISA and ELISpot assays

To detect total Ig by ELISA, 96 well half-area ELISA plates (Corning 96-Well Half Area Flat Bottom Polystyrene High Bind) were coated for 3 h at room temperature with purified goat anti-mouse IgG, IgM, and IgA antibodies (Southern Biotech) used at a concentration of 10 μg/ml in PBS, or with the antigen probes (namely resurfaced eOD-GT8, eOD-GT8-KO2 and HxB2^N276D^), used at a final concentration of 2 ug/ml in PBS. After four washes with PBS 0.025% Tween-20 and blocking with PBS 3% BSA for 1 h at room temperature, samples and standards (relative unlabeled mouse Ig; Southern Biotech) were diluted and incubated at room temperature for 4 h. Specific secondary goat anti-mouse Ig conjugated with horseradish peroxidase (HRP) were added after four washes with PBS and 0.025% Tween-20 and incubated for 2 h at room temperature. Plates were washed again, and the assay was developed with 1-Step™ Ultra TMB-ELISA Substrate Solution (Thermo Fisher, Cat#34029) and blocked with 2N H_2_S0_4_ solution before reading. Plates were read at 450 nm within 1 hour after the stop solution, using an Envision 2104 Multilabel Plate Reader (PerkinElmer).

Antibody secreting cells (ASC) were detected using an ELISPOT assay. Briefly, 96-well plates (Millipore, MSIPS4510 Sterile, hydrophobic high protein binding Immobilon-P membrane) were coated with 10 μg/ml purified goat anti-mouse IgM (Catalog #1021-01), IgG (Catalog #1030-01) or IgA (Catalog #1040-01) (all from Southern Biotech) for the detection of total ASC, or with eOD-GT8 resurfaced monomer probes (WT and KO) for the detection of antigen-specific ASC. Plates were left incubating for 2 h at room temperature. After three washes with PBS solution, plates were blocked with PBS and 1% BSA and incubated for 30 min at 37°C. Serial dilutions of splenocytes or cultured B cells were added in a final volume of 200 μl B cell complete medium and left at 37°C overnight. Subsequently, plates were washed three times with PBS and 0.25% Tween-20 and four times with PBS and incubated for 2 h at room temperature with biotinylated goat anti-mouse IgM (Catalog #1020-08), IgG (Catalog #1030-08) or IgA (Catalog #1040-08) (all from Southern Biotech). After washing, avidin-peroxidase (HRP; Sigma-Aldrich, Catalog #A3151) was added and left for 1 h at room temperature. The assay was developed with AEC (Sigma-Aldrich, Catalog #A6926). For quantification of ASC, plates were acquired, counted, and quality controlled using an ELISPOT reader and ImmunoSpot 5.1 software (CTL).

## QUANTIFICATION AND STATISTICAL ANALYSIS

Statistical analyses were performed using GraphPad Prism v9. Data were statistically significant at * p < 0.05, ** p < 0.01, *** p < 0.001, and **** p < 0.0001.

