## Supplemental figures for "IL-2 secreting T helper cells promote EF B cell maturation via intrinsic regulation of B cell mTOR/AKT/Blimp-1 axis"

### Supplemental Table

| sgRNA | sequence |
|---|---|
| *Aicda1.1 F1* | CACCGCGCTGGAGACCGATATGGAC |
| *Aicda1.1 R1* | AAACGTCCATATCGGTCTCCAGCGC |
| *Bach2.1 F1* | CACCGTTATACCGAACAGAGACCGC |
| *Bach2.1 R1* | AAACGCGGTCTCTGTTCGGTATAAC |
| *Cd8a.1 F1* | CACCGGCAGGTTCAGCGACAGAAAG |
| *Cd8a.1 R1* | AAACCTTTCTGTCGCTGAACCTGCC |
| *Foxo1.1 F1* | CACCGTCCAGTCCGGCGCCGTCGGG |
| *Foxo1.1 R1* | AAACCCCGACGGCGCCGGACTGGAC |
| *HVEM.1 F1* | CACCGCCTGAAGGTGTTGTCTGTAG |
| *HVEM.1 R1* | AAACCTACAGACAACACCTTCAGGC |
| *ICOS.1 F1* | CACCGCTGAAGCTCTGGCTACCCGT |
| *ICOS.1 R1* | AAACACGGGTAGCCAGAGCTTCAGC |
| *IL2.1 F1* | CACCGAAGATGAACTTGGACCTCTG |
| *IL2.1 R1* | AAACCAGAGGTCCAAGTTCATCTTC |
| *IL2Ra.1 F1* | CACCGGGTGCCGTTCTTGTAGGAGA |
| *IL2Ra.1 R1* | AAACTCTCCTACAAGAACGGCACCC |
| *IRF4.1 F1* | CACCG GCAATGGGAAACTCCGACAG |
| *IRF4.1 R1* | AAACCTGTCGGAGTTTCCCATTGC C |
| *prdm1.1 F1* | CACCGAGAAGTACCCCCGTCAGCGC |
| *prdm1.1 R1* | AAACGCGCTGACGGGGGTACTTCTC |
| *tlr7.1 F1* | CACCGACTTAGACTTCTCCAACAAC |
| *tlr7.1 R1* | AAACGTTGTTGGAGAAGTCTAAGTC |

**Table S1.** sgRNA sequences designed with the CHOP-CHOP website. Related to Figures 2, 3, 4, 6 and S4, S6, S7, S8, S11.

### Supplemental Figures

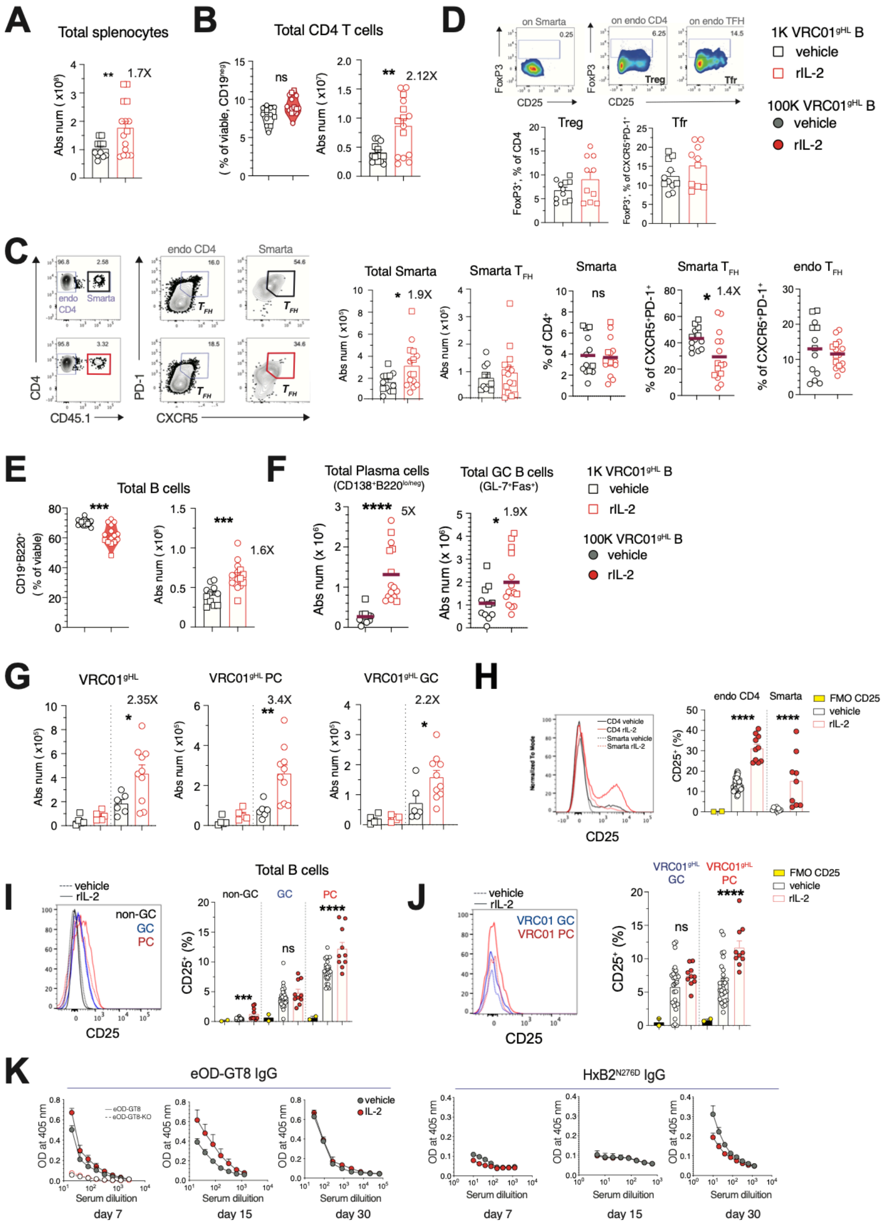


**Figure S1. Effect of in vivo rIL-2 treatment on adaptive immune cells, Related to Figures 1**

Analysis of cell in spleens of vehicle-treated and rIL-2 injected mice on day 7. (A) Quantification of total splenocytes. (B) Quantification of total CD4 T cells. (C) Representative dot plots and quantification of total Smarta cells, Smarta Tfh and endogenous Tfh cells (D) FACS plots and frequencies of FoxP3 expressing cells, Treg and Tfr. (E) Frequency and count of total B cells (F) Absolute count of total PC and GC B cells in the indicated groups of mice. (G) Absolute count of VRC01$^{gHL}$ uGFP B cells and relative populations of PC and GC. (H) Histogram and quantification of CD25 positive cells on endogenous CD4 and Smarta T cells. (I) Histogram and quantification of CD25 positive endogenous B cells (J) Histogram and quantification of CD25 expressing VRC01$^{gHL}$ PC and GC subsets. The FMO is showing the CD25 negative signal on each population. (K) ELISA serology for eOD-GT8 and HxB2(N276D) probes at each indicated time-points. Data are combined from all control or rIL-2 treated mice receiving low pf (1K, squares) or high pf (100K, circles) of VRC01$^{gHL}$ uGFP. Each dot represents an individual mouse; data are from n=4 pooled experiments. P values were determined by two-tailed Student's t test. Not significant, ns $p > 0.05$; *$p < 0.05$; **$p < 0.01$; ***$p < 0.001$; ****$p < 0.0001$

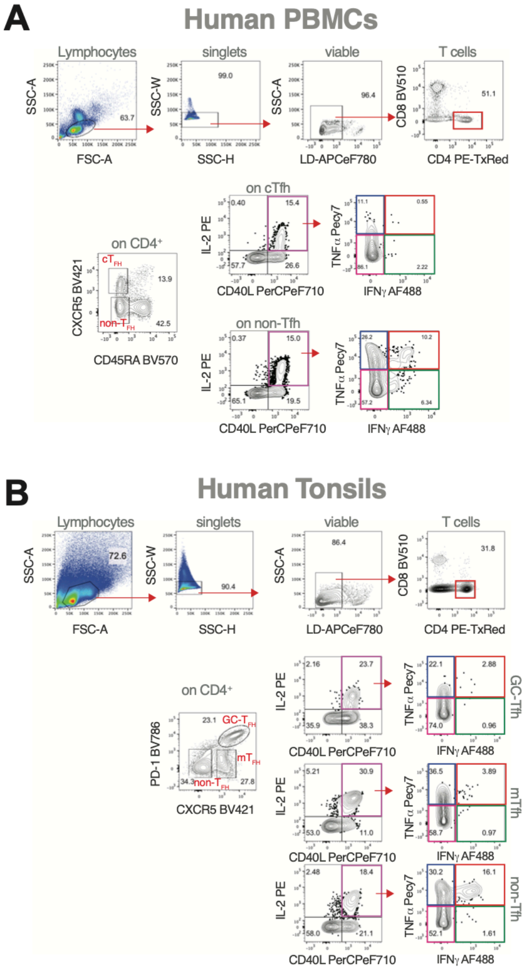


Figure S2. ICS assay to detect IL-2 secreting antigen-specific CD4 T cells isolated from PBMCs and tonsils, Related to Figures 2

Gating strategy to quantify IL-2 secreting CD40L expressing CD4 T cells and co-secretion of TNF-a and IFN-g cytokines in (A) PBMCs or (B) tonsils from healthy donors. The ICS assay was performed after 6h of stimulation *in vitro*.

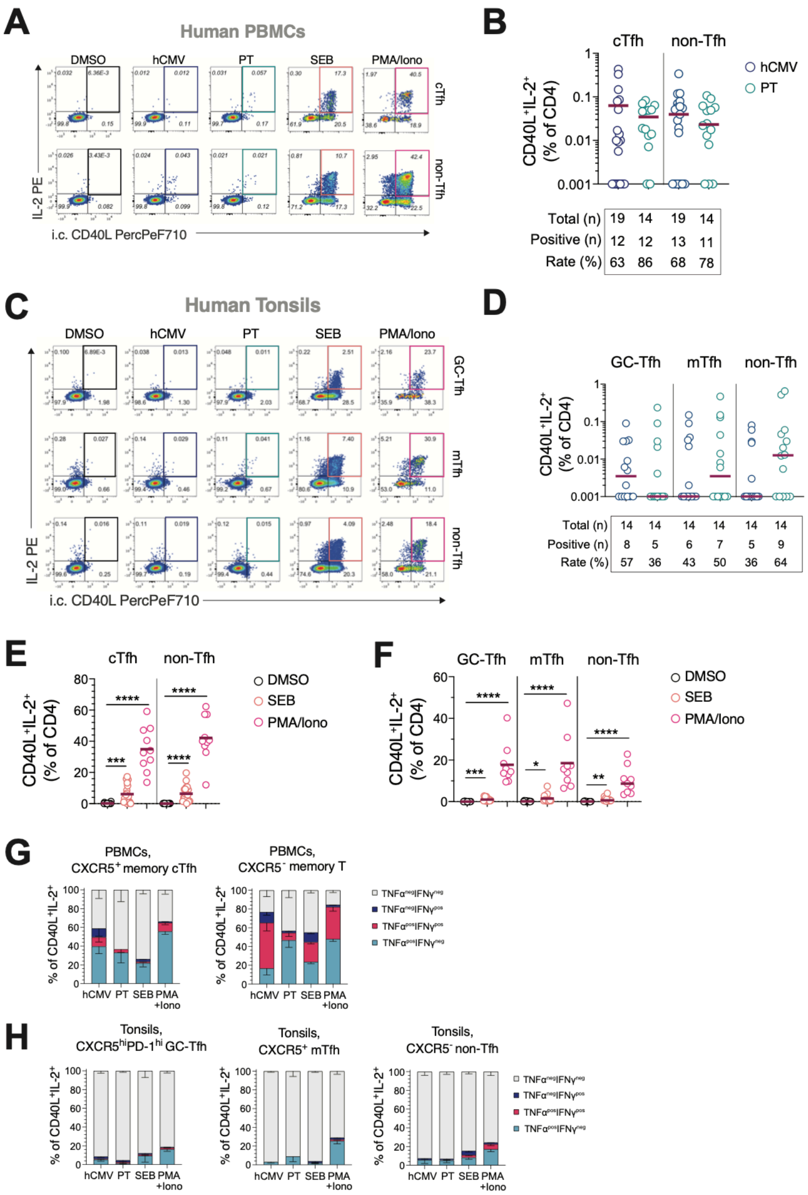


**Figure S3. Characterization and quantification of antigen-specific IL-2$^+$ T helper cells in blood and tonsils, Related to Figure 2**

(A) FACS plots representing the gating strategy to assess IL-2 secreting CD40L expressing T cells in PBMC and (B) quantification of their frequency on total CD4 upon stimulation with hCMV or PT mega pools of peptide (MPs). (C) FACS plots representing the gating strategy to assess IL-2 secreting CD40L expressing T cells in tonsils and (D) relative quantification upon stimulation with human Cytomegalovirus (hCMV) or pertussis (PT) mega pools of peptide (MPs) (E) CD40L$^+$IL-2$^+$ induced by SEB or PMA/ionomycin stimulations of PBMCs. (F) CD40L$^+$IL-2$^+$ induced by SEB or PMA/ionomycin stimulations of tonsils. Co-secretion of TNF-a and IFN--g in response to the indicated stimulations quantified for all the indicated CD40L$^+$IL-2$^+$ expressing T helper subsets in blood (G) or (H) tonsils. Each dot represents an individual donor. Data are combined from n=4 experiments for PBMCs (n= 10-20 subjects) or n= 3 Tonsils (n= 9-14 subjects) samples. Line represents the mean. P values were determined by paired one-way ANOVA test. Not significant, ns $p > 0.05$; *$p < 0.05$; **$p < 0.01$; ***$p < 0.001$; ****$p < 0.0001$

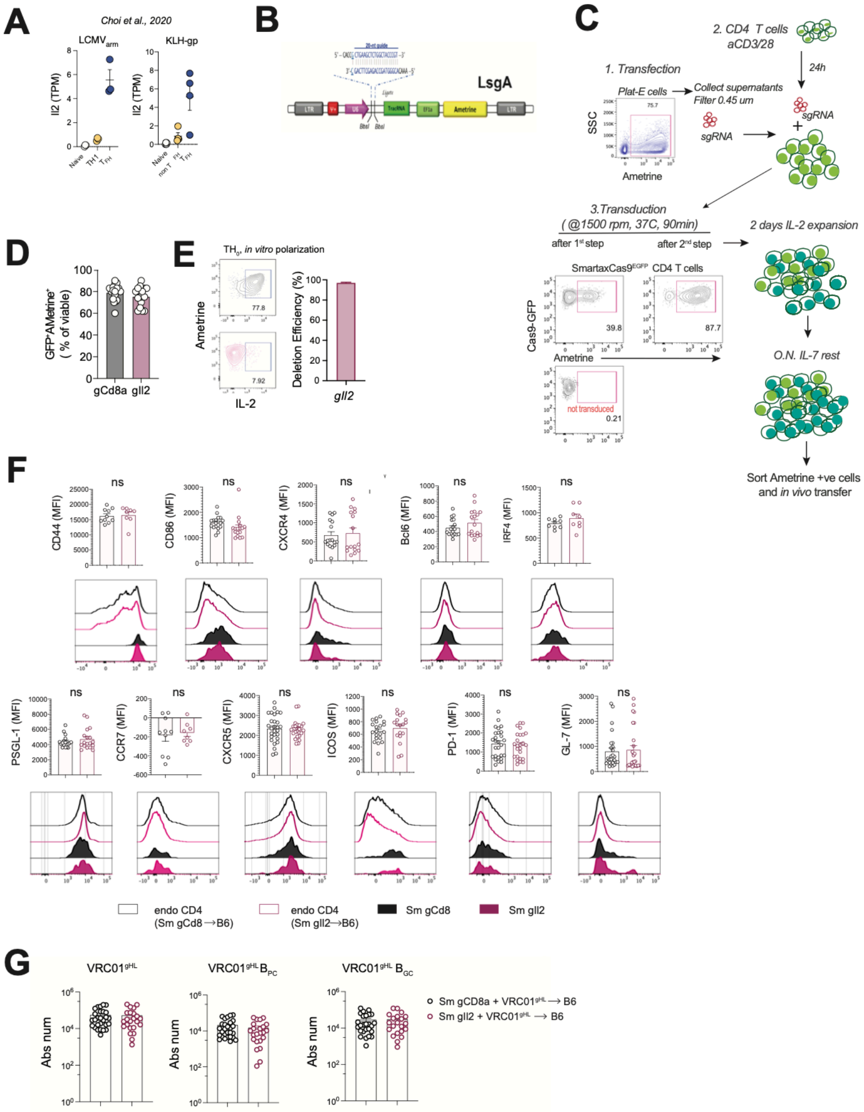


**Figure S4. CRISPR/Cas9 mediated generation of gIl2 (IL-2$^{-/-}$) Smarta T cells, Related to Figure 2**

(A) RNA-seq data showing Il2 transcripts expression (TPM) in sorted Smarta cells subsets from day 7 of LCMV-Armstrong (LCMV-Arm) infection or gp-KLH/Alum (KLH-gp) immunization (data are from Choi et al., 2020). (B) Scheme of the LsgA plasmid used to deliver sgRNA into activated cells with retroviral transduction protocols. (C) Schematic for activation and retroviral transduction of murine CD4 T cells. (D) Quantification of retroviral transduction efficiency on Smarta CD4 T cells assessed after a 2-steps transduction. gCd8 and gIl2 shown as a percentage of viable, GFP (Cas9$^{EGFP}$) and Ametrine (LsgA vector reporter) positive cells. (E) Validation of gIl2 efficiency in disrupting IL-2 locus assessed by testing the IL-2 secretion *in vitro* upon forced Th-0 conditions (10 µg/mL anti-IFN-g, anti-IL-4, and anti-TGF-b (Eto et al., PloS One 2011)) of gCd8 Smarta cells. (F) Analysis of gCd8 and gIl2 Smarta cells one week after eOD-GT5$_{gp61}$60mer immunization. MFIs comparisons are relative to the expression of activation markers and TFs associated with T cells activation, differentiation, and migration. Histograms show endogenous CD4 T cells from recipient mice for reference. Data are combined from n= 3 or 4 experiments. Each dot represents an individual mouse. P values were determined by a two-tailed Student's t-test. Not significant, ns $p > 0.05$; *$p < 0.05$; **$p < 0.01$; ***$p < 0.001$; ****$p < 0.0001$.

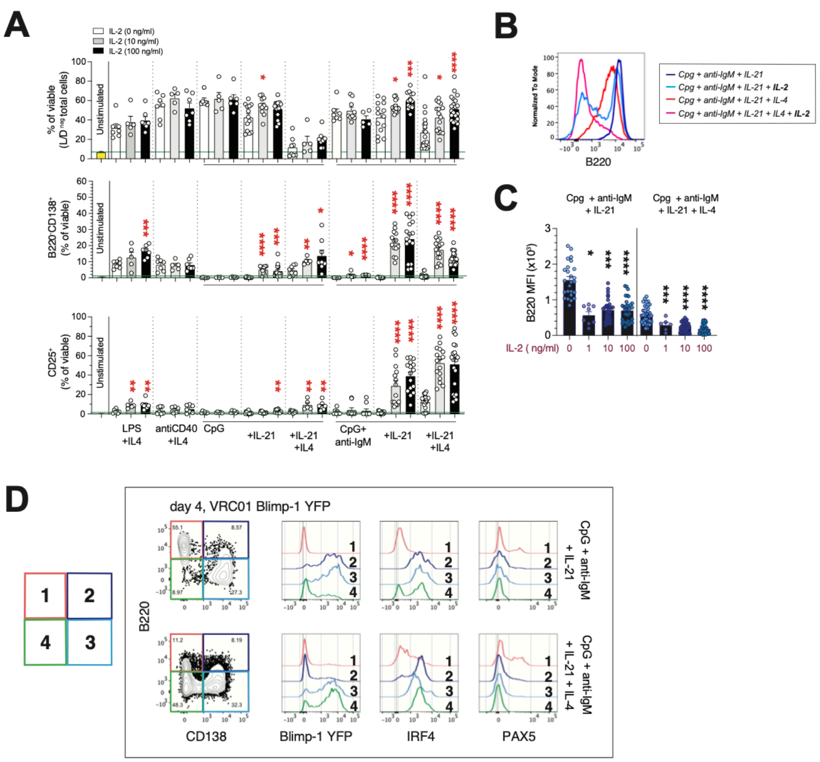


**Figure S5.** *In vitro* **culture system used to generate terminally differentiated murine plasma cells, Related to Figures 3, 4 and 5**

(A) Comparison of viability, plasma cell frequency, and CD25 expression on naïve murine B cells stimulated for 4 days with the indicated cocktails of stimuli. (B) Downregulation of B220 on activated B cells (day 4) is shown and quantified for each indicated condition. (C) Expression of Blimp-1, IRF-4, and PAX-5 on VRC01$^{gHL}$ blimp-1 YFP-reporter cells stimulated for four days with each indicated cocktail of stimuli. Subsets of viable activated B cells, at different stages of differentiation, are shown in red (#1, B220$^{+}$CD138$^{-}$), dark blue (#2, B220$^{+}$CD138$^{+}$), light blue (#3, B220$^{-}$CD138$^{+}$) and green (#4, B220$^{-}$CD138$^{-}$). Data are from n=13 pooled individual experiments. P values were determined by a two-tailed Student's t-test. Not significant, ns $p > 0.05$; *$p < 0.05$; **$p < 0.01$; ***$p < 0.001$; ****$p < 0.0001$.

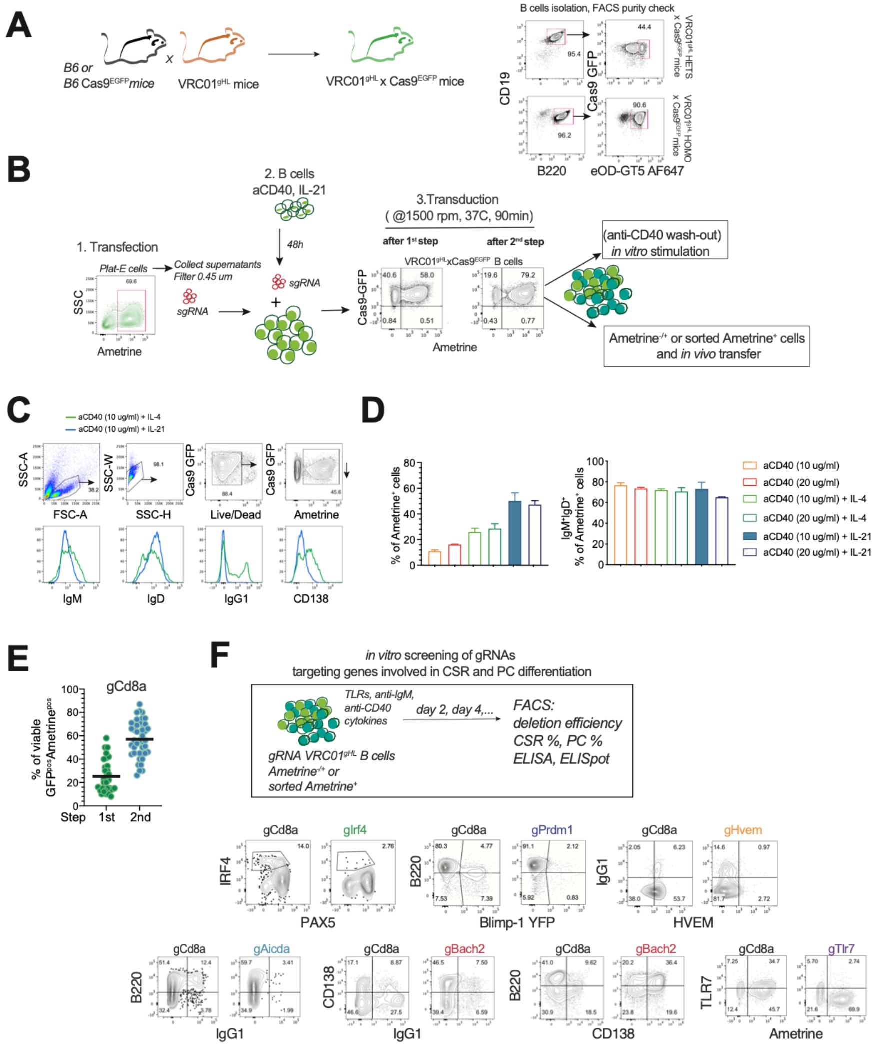


**Figure S6. A novel CRISPR/Cas9 B cell targeting approach to generate knock-out B cells, Related to Figures 3, 4 and 6**

(A) Generation of VRC01$^{gHL}$xCas9$^{EGFP}$ mice and validation of B cells binding to eOD-GT5 AF647 probe from heterozygous or homozygous mice. (B) Schematic for the activation and retroviral transduction of murine naïve B cells and relative analysis performed in the study. (C) Validation of the novel protocol for retroviral transduction of murine B cells tested with different combinations and doses of anti-CD40 and cytokines (green line rIL-4, blue line rIL21). Dot plots showing the expression of IgM and IgD, IgG1 and CD138 on Ametrine$^{+}$ transduced B cells after a 2-steps retroviral transduction. (D) Quantification of Ametrine$^{+}$ transduced B cells and percentage of IgM, IgD double expressing naïve-like transduced B cells after a 2-steps retroviral transduction is shown for each indicated combination of stimuli. Data are from n=4 experiments. Error bars represent the SEM. (E) Quantification of retroviral transduction efficiency of gCD8 VRC01$^{gHL}$xCas9$^{EGFP}$ B cells assessed after each transduction step. Data are from n=26 pooled experiments. Each dot represents an individual gRNA tested sample. (F) Schematic representation of gRNA screening *in vitro* for testing the efficiency of gene disruptions and relative B cells differentiations. Representative FACS plots showing well-known target genes and their effect on B cells. (G) Quantification of retroviral transduction efficiency on gIl2ra VRC01$^{gHL}$xCas9$^{EGFP}$ B cells assessed after each transduction step. Data are from n=14 pooled experiments. Each dot represents an individual gRNA tested sample. (H) Histogram showing the expression of CD25 after targeting transduction of B cells with gCd8 (control) and gIl2ra (CD25 disruption) and stimulation with the indicated combination of stimuli. Data are from n=10 experiments analyzed after 2 days post-stimulation. (I) Efficiency of deletion of CD25 expression on stimulated gIl2ra B cells (as in H). Data are from n=10 pooled experiments. Each dot represents an individual gRNA tested sample. (J) Schematic of Electroporation (EP) of crRNA ATTO-550 PE delivered to murine activate B cells with RNP complex. (K) Histogram showing the detection of ATTO-550 in the PE channel with crCd8 (control) and crIl2ra (CD25 KO) guides, in comparison to electroporated B cells without RNP delivery. (L) Quantification of the frequency of ATTO-550 PE electroporated B cells calculated 24 hours post-EP for the indicated crRNA guides. (M) Representative FACS plots showing the expression of B220, CD138 and CD25 on and (N) quantification of viable, CD25 and CD138 positive cells after stimulation with CpG, rIL-21 and rIL-2 (day 4) for the indicated crRNA guides. Data are from n=4 pooled experiments. Each dot represents an individual gRNA tested sample. P values were determined by a two-tailed Student's t-test. Not significant, ns $p > 0.05$; *$p < 0.05$; **$p < 0.01$; ***$p < 0.001$; ****$p < 0.0001$.

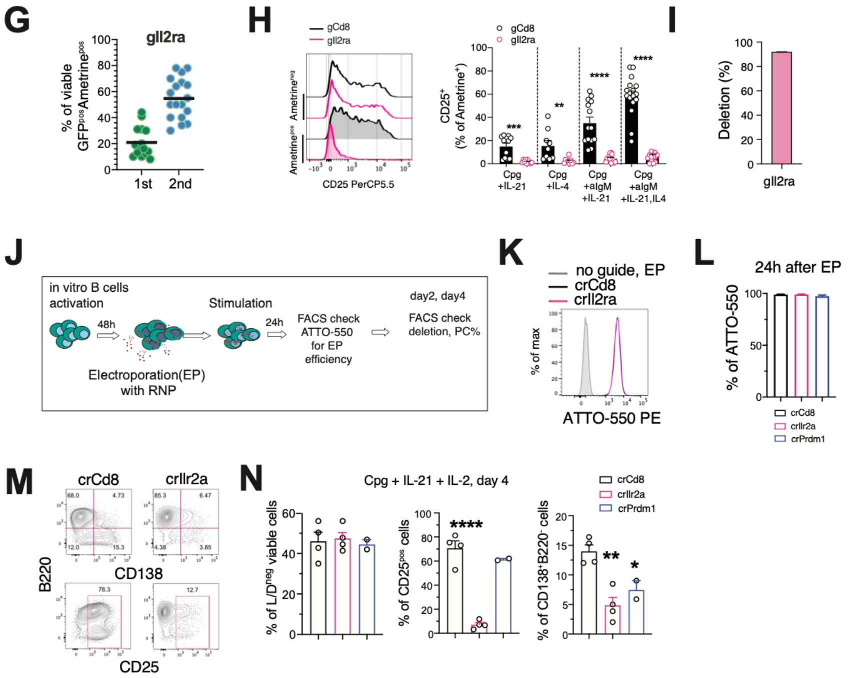


Figure S6. A novel CRISPR/Cas9 B cell targeting approach to generate knock-out B cells, Related to Figures 3, 4 and 6 *(continue)*

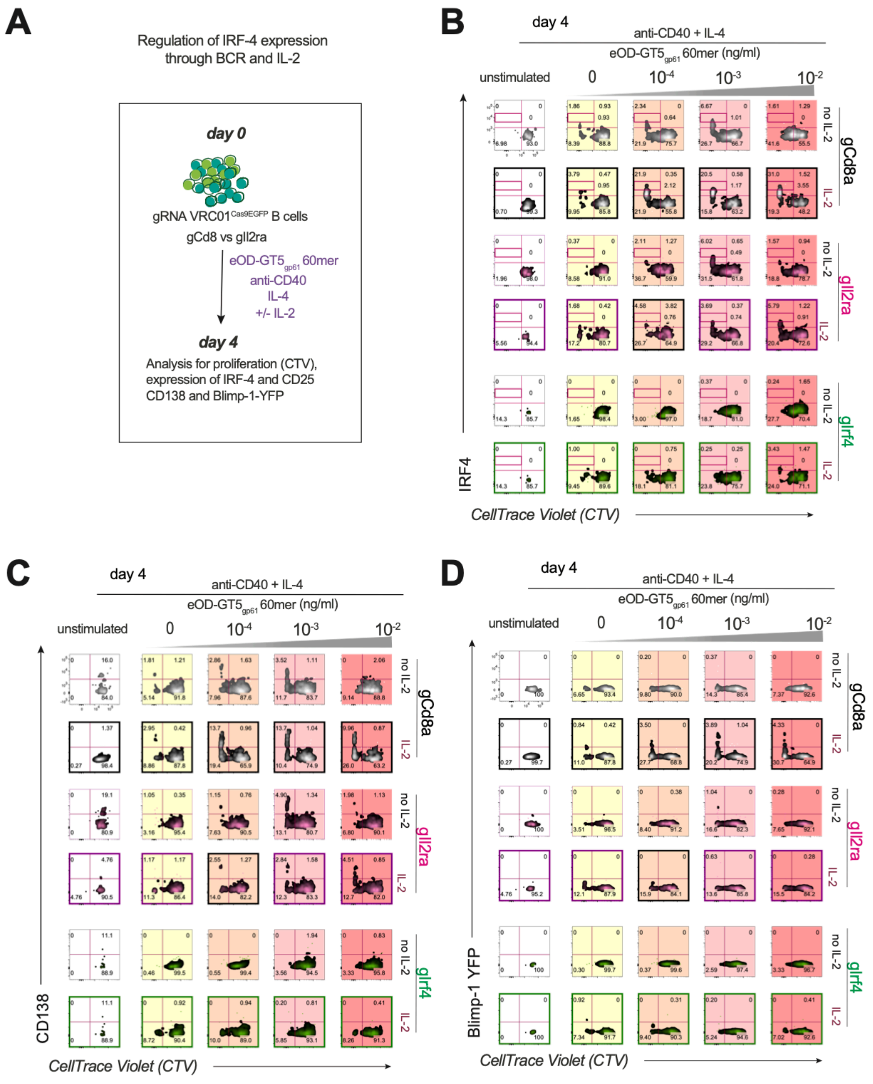


**Figure S7. Regulation of IRF4 expression through BCR and IL-2 signals in cultured B cells, Related to Figure 4**

(A) Schematic diagram for the analysis *in vitro* of the regulation of IRF4 on gRNA VRC01$^{gHL}$xCas9$^{EGFP}$ transduced stimulated B cells. Testable model for IRF4$^{hi}$ expressing plasma cells in the presence of increasing BCR signaling. (B) Representative FACS plots for day 4 analysis of transduced gCd8, gIl2ra or gIrf4 VRC01$^{gHL}$xCas9$^{EGFP}$ treated without or with IL-2 (100 ng/ml). Populations of CTV$^{lo}$, IRF4$^{pos}$ and IRF4$^{high}$ proliferated B cells are gated on viable, Ametrine positive (transduced) cells. (C) Representative FACS plots showing CTV$^{lo}$, CD138$^{pos}$ and (D) CTV$^{lo}$, Blimp-1$^{pos}$ cells on day 4. Data are from n=3 independent experiments. P values were determined by a two-tailed Student's t-test. Not significant, ns $p > 0.05$; *$p < 0.05$; **$p < 0.01$; ***$p < 0.001$; ****$p < 0.0001$.

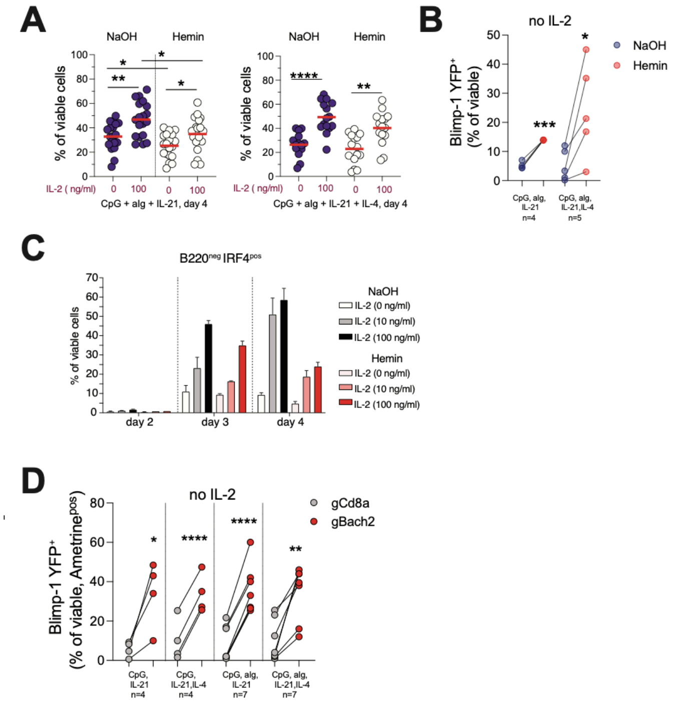


**Figure S8. IL-2 and BACH2 regulation of prdm1/Blimp-1 expression on *in vitro* stimulated B cells, Related to Figure 4**

(A) Viable cells quantification upon NaOH or Hemin treatment of naïve B cells stimulated for 4 days *in vitro*. (B) Blimp-1 quantification of cells treated as in (A). Data are from n=4 experiments (C) Quantification of B220$^{lo/neg}$IRF4$^{pos}$ B cells from day 2 to day 4 upon NaOH or Hemin treatment of naïve B cells. Each dot represents an individual experiment. Data are from n=4 combined experiments. (D) Blimp-1 quantification of gRNA transduced control or BACH2 KO VRC01$^{gHL}$ B cells in the absence of IL-2. Each dot represents an individual experiment. Data are from n=4 combined experiments. P values were determined by a two-tailed Student's t-test. Not significant, ns $p > 0.05$; *$p < 0.05$; **$p < 0.01$; ***$p < 0.001$; ****$p < 0.0001$.

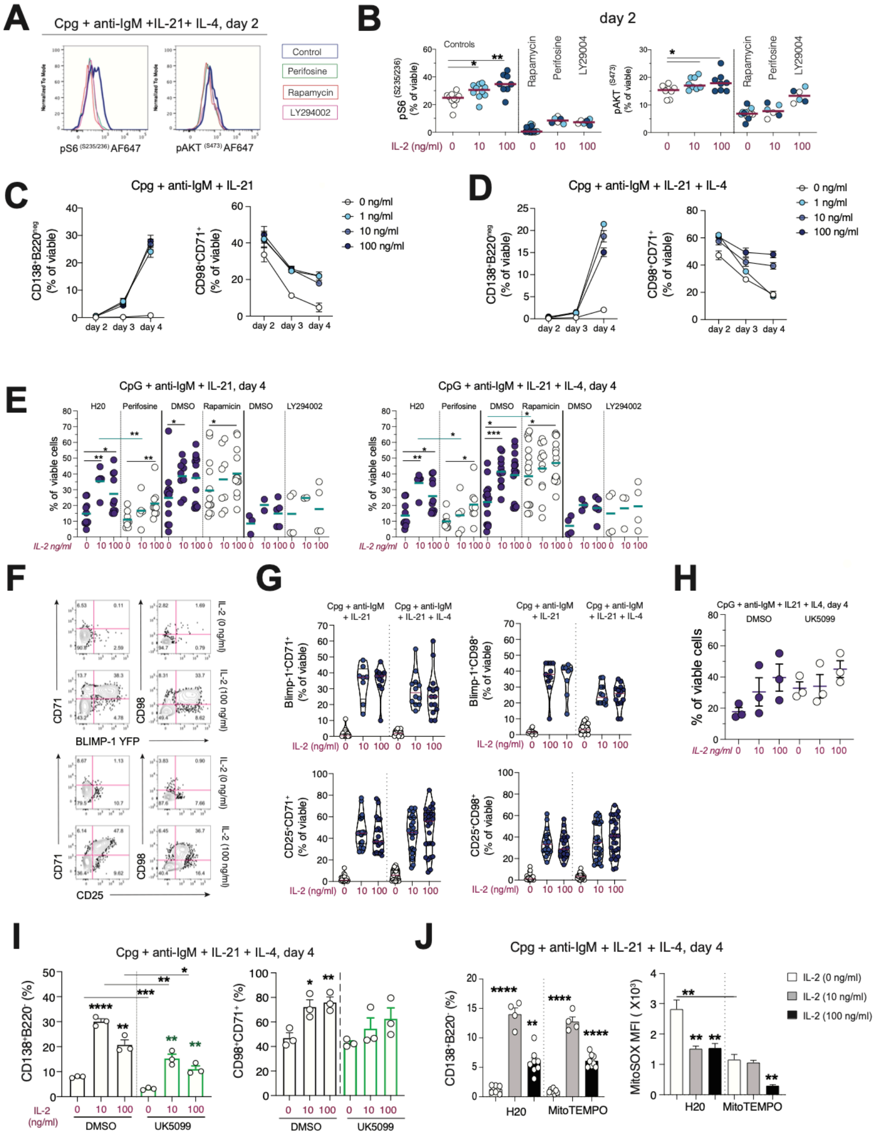


**Figure S9. IL-2/CD25 metabolic regulation of B cells differentiation through mTOR and Blimp-1 axis, Related to Figure 5**

(A) Histograms showing the expression of phosphorylated S6 (pS6$^{(S235/236)}$ AF647 and phosphorylated AKT (pAKT$^{(S473)}$ AF647) in stimulated B cells treated with Perifosine (green line), Rapamycin (red line) or LY294002 (purple line) inhibitors and compared to control cells (blue line). (B) Relative quantification after 2 days of culture of cells treated as in (A). Data are from n=2 independent experiments. Each dot represents an individual sample tested. Kinetics of plasma cells (B220$^{-}$CD138$^{+}$) and CD98, CD71 expression on days 2, 3 and 4 post- stimulation of naïve B cells with (C) CpG + anti-IgM and IL-21 or (D) CpG + anti-IgM and IL-21+ IL-4. (E) Viability of B cells calculated upon each indicated drug tested and assessed on day 4 for each stimulation as indicated. Data are from n=6-7 combined experiments. (F) FACS plots representative of the expression of CD71, CD98 on Blimp-1 YFP expressing B cells, and CD71 and CD98 on CD25 expressing B cells. (G) Relative quantifications for each subset and indicated conditions after 4 days of culture. Data are from n=5 independent experiments. Each dot represents an individual sample. (H) Effect of UK5099 (mitochondrial pyruvate transporter inhibitor) on B cells viability, and (I) plasma cells formation and CD98, CD71 expression after 4 days of B cells culture. Data are from n=2 independent experiments. Each dot represents an individual well. (J) Plasma cell generation and ROS accumulation (MitoSOX staining) in control (H2O) and MitoTEMPO treated B cells, at different doses of rIL-2, after 4 days of stimulation. Data are from n=4 independent experiments. Each dot represents an individual well. P values were determined by a two-tailed Student's t-test. Not significant, ns $p > 0.05$; *$p < 0.05$; **$p < 0.01$; ***$p < 0.001$; ****$p < 0.0001$.

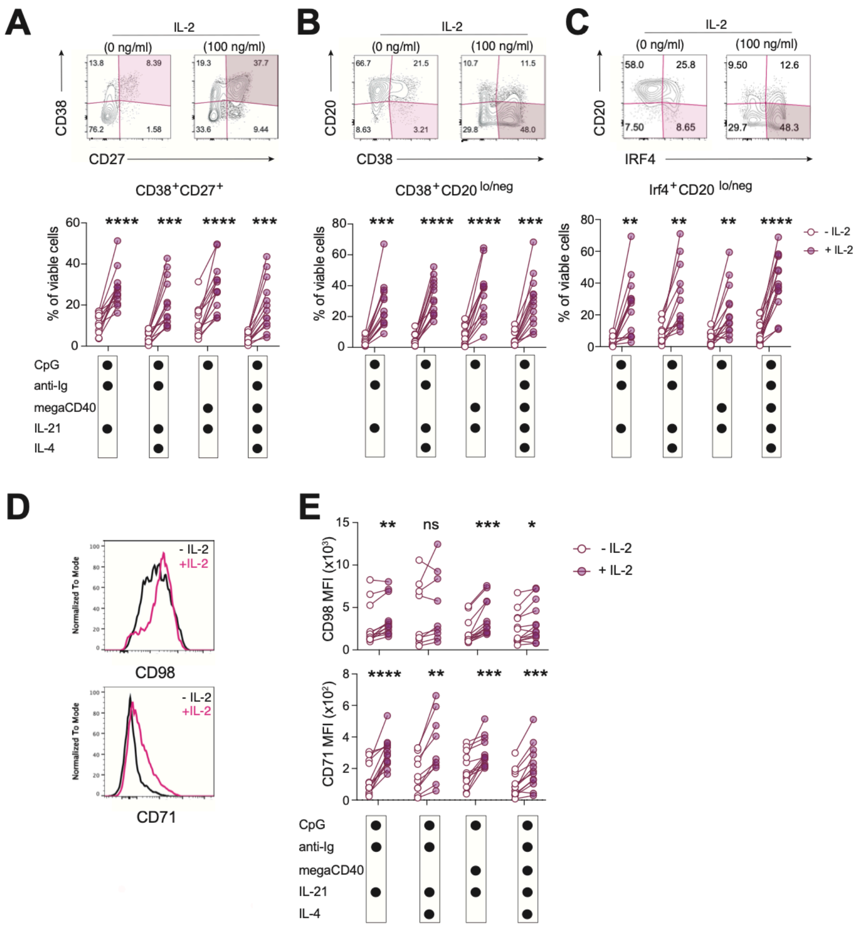


**Figure S10.** ***In vitro*** **culture system to generate terminally differentiated human plasma cells and their metabolic regulation via IL-2 signals, Related to Figure 5**

Representative dot plots and relative quantification of viable B cells expressing (A) CD38$^{+/hi}$ CD27$^{+/hi}$, (B) CD20$^{lo/-}$ CD38$^{+/hi}$ and (C) CD20 $^{lo/-}$ IRF4$^{+}$ to quantify *in vitro* generated human plasma cells. Isolated naïve B cells from HD (n=13) were stimulated for 7 days in the presence of the indicated stimuli (black-filled circles in the graphs). (D) Representative histograms showing the upregulation of CD98 and CD71 in the presence of IL-2, and (E) relative MFIs quantification for each cocktail of stimuli tested, as indicated by the black-filled circles in the graphs. Data are from n=3 independent experiments. P values were determined by paired one-way ANOVA test. Not significant, ns $p > 0.05$; *$p < 0.05$; **$p < 0.01$; ***$p < 0.001$; ****$p < 0.0001$.

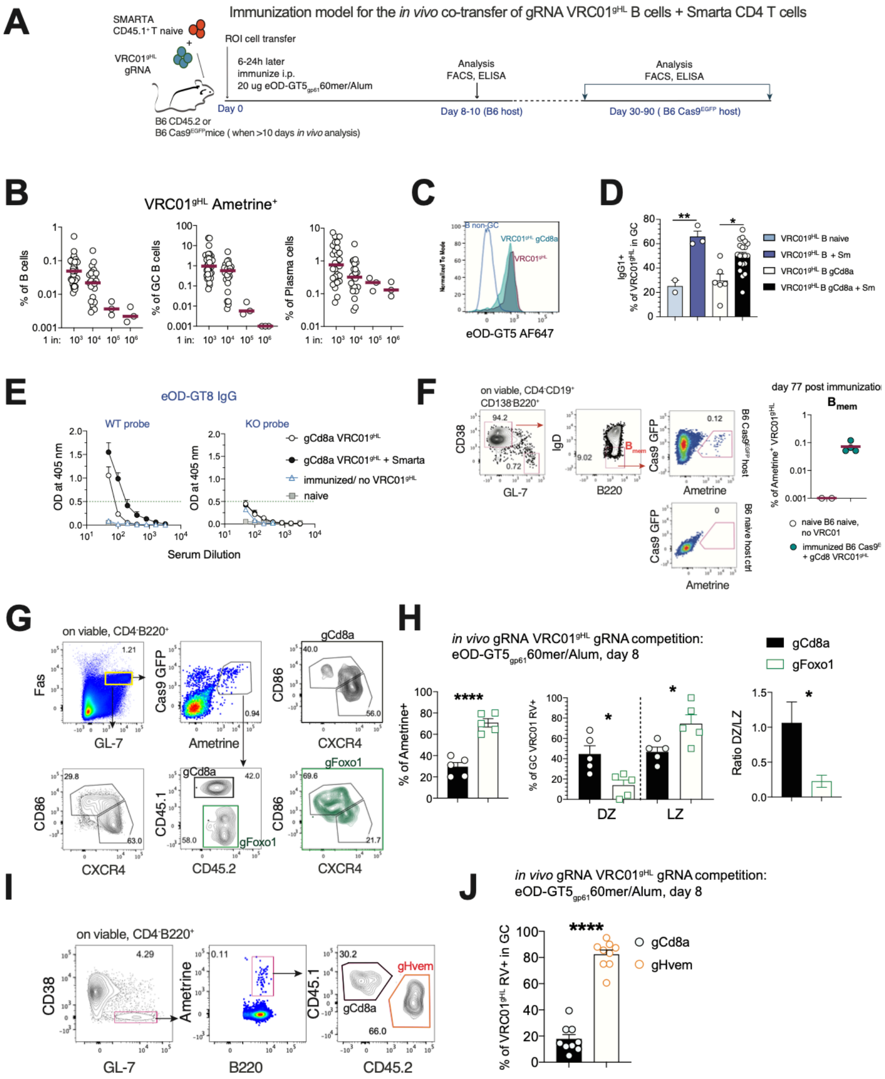


**Figure S11. Novel in vivo transfer system of gRNA VRC01$^{gHL}$xCas9$^{EGFP}$ B cells: validation and analysis of immune responses, Related to Figure 6**

Establishment of a model to study the role of specific genes of interest upon *in vivo* transfer of gRNA transduced VRC01$^{gHL}$xCas9$^{EGFP}$ B cells into recipient B6 mice. (A) Schematic showing the model of *in vivo* transfer of antigen-specific T and B cells. Kinetics of immunization and time-points of analysis are indicated in the scheme. (B) Quantification of Ametrine positive cells showed as a percentage of total B cells, $B_{GC}$ or $B_{PC}$ upon transfer of different numbers of transduced B cells and analyzed on day 8-10 post-immunization to assess PC and GC B cells differentiation in the spleen. (C) Histogram showing the relative binding to the eOD-GT5 AF647 probe for comparison of transduced gCd8 VRC01$^{gHL}$xCas9$^{EGFP}$, naïve transferred VRC01$^{gHL}$, and endogenous GL-7$^{neg}$Fas$^{neg}$ non-GC B cells, on day 8. (D) Quantification of class switched IgG1 expressing cells from the groups of mice recipient of naïve VRC01$^{gHL}$ or gCd8 transduced VRC01$^{gHL}$xCas9$^{EGFP}$, without or with Smarta T cells transfer (day 8 post-immunization). Each dot represents an individual mouse. (E) Serological evaluation of secreted eOD-GT8 reactive IgG tested in ELISA (WT and KO GT8 probes). Data show the comparison of gCd8 transduced VRC01$^{gHL}$xCas9$^{EGFP}$ without or with Smarta T cells. Sera from naïve (not immunized) and immunized mice (without VRC01$^{gHL}$ transfer) are used as a reference, negative control of the assay. Data are from n=4-9 experiments. (F) Representative dot plots to assess the generation on memory cells from transferred gCd8 VRC01$^{gHL}$xCas9$^{EGFP}$ B cells and the relative quantification of viable, CD4$^-$CD19$^+$CD138$^-$B220$^+$GL-7$^-$CD38$^+$IgD$^-$ Ametrine positive memory cells analyzed 77 days after immunization. Data are from n=2 experiments. *In vivo* gRNA VRC01$^{gHL}$xCas9$^{EGFP}$ competition model: gRNAs VRC01$^{gHL}$xCas9$^{EGFP}$ are co-transferred into wild-type recipient mice and spleens are analyzed after 8 days. (G) Representative dot plots and (H) quantification of gCd8 and gFoxo1 VRC01$^{gHL}$xCas9$^{EGFP}$ transduced B cells differentiated into DZ (CXCR4$^+$CD86$^-$) and LZ (CXCR4$^-$CD86$^+$) GC B cells. Data are from n=2 experiments. (I) Representative dot plots and (J) quantification of gCd8 and gHvem VRC01$^{gHL}$xCas9$^{EGFP}$ transduced B cells showed as a percentage of total $B_{GC}$ cells. Data are from n=4 experiments. Error bars represent the SEM. P values were determined by a two-tailed Student's t-test. (K) FACS plots and (L) quantification of total Ametrine$^{pos}$ transferred cells and $B_{PC}$ generated with gPrdm1 VRC01 $^{gHL}$ transduced B cells. (M) Absolute counts of VRC01$^{gHL}$ B cells transferred into recipient immunized mice and analyzed on day 8, for each indicated guide. Data are from n=6 experiment for gCD8 (34 mice), n=4 experiments for gIl2ra (20 mice), n=4 experiments for gPrdm1 (n=13 mice) and n=2 experiments for gAicda (n=9 mice). (N) MFIs and quantification of IgG1 expressing cells from the indicated transduced VRC01$^{gHL}$ B cells. Each dot represents an individual mouse. Error bars represent the SEM. P values were determined by a two-tailed Student's t-test. Not significant, ns $p > 0.05$; *$p < 0.05$; **$p < 0.01$; ***$p < 0.001$; ****$p < 0.0001$.

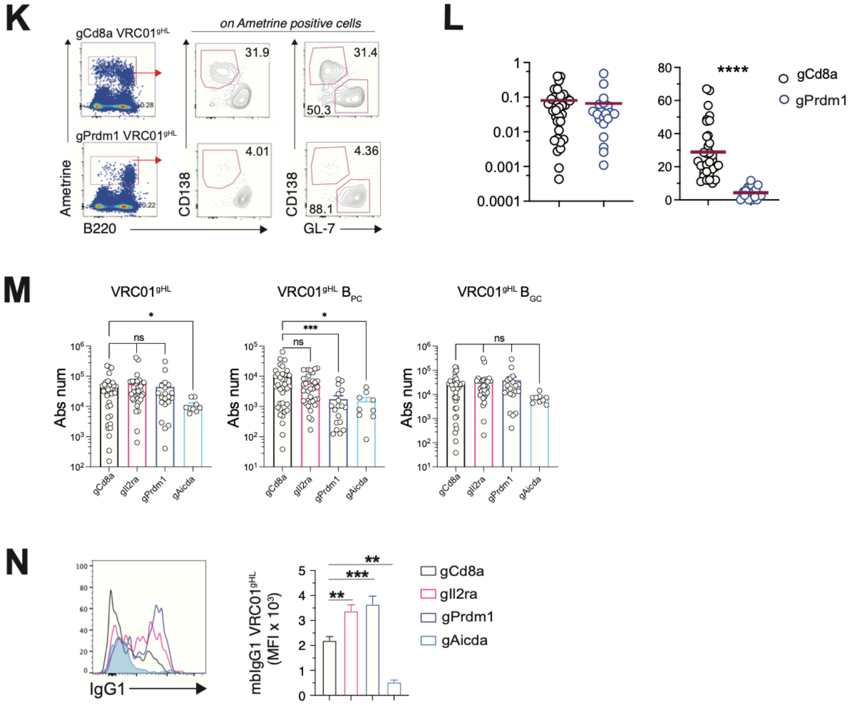


Figure S11. Novel in vivo transfer system of gRNA VRC01$^{gHL}$xCas9$^{EGFP}$ B cells: validation and analysis of immune responses, Related to Figure 6
(continue)
